# Biased Signaling in Mutated Variants of *β*_2_-Adrenergic Receptor: Insights from Molecular Dynamics Simulations

**DOI:** 10.1101/2023.09.14.557674

**Authors:** Midhun K. Madhu, Kunal Shewani, Rajesh K. Murarka

## Abstract

The molecular basis of receptor bias in G protein-coupled receptors (GPCRs) caused by mutations that preferentially activate specific intracellular transducers over others remains poorly understood. Two experimentally identified biased variants of *β*_2_-adrenergic receptors (*β*_2_AR), a prototypical GPCR, are a triple mutant (T68F, Y132A, and Y219A) and a single mutant (Y219A); the former bias the receptor towards the *β*-arrestin pathway by disfavoring G protein engagement, while the latter induces G protein signaling explicitly due to selection against GPCR kinases (GRKs) that phosphorylate the receptor as a prerequisite of *β*-arrestin binding. Though rigorous characterizations have revealed functional implications of these mutations, the atomistic origin of the observed transducer selectivity is not clear. In this study, we investigate the allosteric mechanism of receptor bias in *β*_2_AR using microseconds of all-atom Gaussian accelerated molecular dynamics (GaMD) simulations. Our observations reveal distinct rearrangements in transmembrane helices, intracellular loop 3, and critical residues R131^3.50^ and Y326^7.53^ in the conserved motifs D(E)RY and NPxxY for the mutant receptors, leading to their specific transducer interactions. The reorganization of allosteric communications from the extracellular agonist BI-167107 to the intracellular receptor-transducer interfaces drives the conformational rearrangements responsible for receptor bias in the single and triple mutants. The molecular insights into receptor bias of *β*_2_AR presented here could improve the understanding of biased signaling in GPCRs, potentially opening new avenues for designing novel therapeutics with fewer side effects and superior efficacy.

## Introduction

G protein-coupled receptors (GPCRs) are the largest family of transmembrane proteins that transmit signals from extracellular space to the cell interior.^1^ They regulate almost every physiological and pathophysiological process in humans and are the target of over one-third of the currently marketed drugs for various diseases such as inflammation, central nervous system disorders, cardiovascular and metabolic diseases, and cancer.^2–6^ In GPCR pathways, three classes of cytosolic proteins bind with the receptor and act as downstream signal transducers: heterotrimeric G proteins, GPCR kinases (GRK), and *β*-arrestins. A typical GPCR activation process is initiated by the binding of an agonist at the extracellular orthosteric pocket of the receptor, which is allosterically communicated to its cytosolic part that results in the opening of an intracellular cavity, where the heterotrimeric G protein binds.^7–9^ Subsequently, GRKs phosphorylate the carboxy-terminal tail (C-tail) of the activated receptor, thereby enabling the engagement of *β*-arrestin that sterically occludes G protein coupling.^10^ *β*-arrestins further initiate signaling cascades independent of G proteins downstream to the receptor. In pharmacologically targeting GPCRs, uniform signal transduction through both G proteins and *β*-arrestins induced by balance agonists is reported to cause unwanted side effects.^11–13^ Therefore, biased agonists that selectively activate clinically relevant pathways over others have gained widespread attention recently.^14,15^ Nonetheless, due to a limited understanding of the fundamental principles underlying the mechanism of biased signaling, the development of transducer-selective drugs for GPCRs remains a formidable challenge.^15,16^

In addition to agonists, mutations of critical residues have also been described to elicit distinct functional outcomes (receptor bias), which has helped identify the reorganized residue communication pathways in GPCRs that give rise to biased signaling.^17–23^ The growing evidence from structural and spectroscopic studies suggests that biased agonists and biasing mutations induce specific downstream signaling pathways by stabilizing distinct receptor conformation; however, the molecular determinants of biased signaling remain poorly understood.^12,13,16,24^ Structural studies provide only static pictures of these highly dynamic receptors, whereas spectroscopic approaches are mostly limited to a localized description of the biased conformational variations. To this effect, atomistic molecular dynamics (MD) simulations offer a powerful complementary tool for exploring ensembles of different functional conformations leading to transducer-specific receptor signaling.^25–28^ Multiple simulation studies have focused on understanding the molecular basis of ligand-directed conformational transitions in GPCRs;^26,29–38^ however, studies on biased mutations (i.e., the receptor bias) have been limited.^19,39–41^

*β*_2_-adrenergic receptor (*β*_2_AR), a prototypical GPCR well known for its druggability in cardiovascular diseases,^42^ is extensively studied for biased signaling using experiments and computer simulations.^17,18,27,33,37,43–46^ A recent MD simulation study has investigated the activation of *β*_2_AR by agonists having diverse transducer selectivities and identified key conformational changes and residues involved in ligand-dependent biased signaling.^33^ *β*_2_AR was also subjected to investigate the effect of a mutation (Y199A) at the extracellular tip of transmembrane helix 5 (TM5) that conferred G protein bias to the receptor.^19^ Nonetheless, the impact of mutations in the selective activation of different transducers of *β*_2_AR is still unexplored. Two experimentally studied biased mutants of *β*_2_AR are (i) a triple mutant T68^2.39^F, Y132^3.51^G, and Y219^5.58^A that induces *β*-arrestin-biased signaling, rationally designed based on evolutionary trace analysis,^17^ and (ii) a single mutant Y219^5.58^A, which causes bias towards G_s_ protein signaling identified using a directed mutagenesis strategy around the same three residues as in the triple mutant^18^ (the superscripts of residues are the Ballesteros–Weinstein numbering scheme of GPCRs^47^). The *β*-arrestin bias of the triple mutant is directly caused by an inefficient coupling of the G_s_ with the receptor.^17^ In contrast, the G protein bias of the single mutant is due to a deficiency in GRK binding that resulted in a lack of C-terminal phosphorylation and subsequent failure of *β*-arrestin engagement with the receptor.^18^ This suggests that the signaling profiles induced by biased ligands downstream to GPCRs can no longer be categorized into the binary classes of G protein-biased and *β*-arrestin-biased, as the interactions with GRKs also regulate the transducer-selectivity of receptors. Therefore, an atomistic understanding of the complex network of interactions between receptors and transducers is essential for the rational design of functionally selective GPCR-targeting drugs. Although the functional outcomes of the single and triple mutants of *β*_2_AR were characterized using experiments, the atomistic basis that governs the structural and dynamic origin of their biased signaling remains to be elucidated.

In this study, we performed large-scale all-atom MD simulations with Gaussian accelerated enhanced sampling (GaMD)^48–50^ to investigate the atomistic basis of receptor bias in *β*_2_AR induced by the single and triple mutations. We extensively compared the conformational ensembles sampled by the mutants with the wild-type, and found that the transmembrane region and intracellular loop 3 (ICL3) undergo distinct rearrangements while conferring the differential bias to the receptor variants. Additionally, suboptimal paths from the extracellular agonist to the intracellular transducer interfaces in the mutant systems show alterations in allosteric communications, leading to transducer-specific downstream signaling.

## Materials and Methods

### System preparations and molecular dynamics (MD) simulations

After removing the co-crystallized G_s_, camelid antibody fragment, and T4 lysozyme from the active crystallographic structure (PDB ID: 3SN6),^51^ *β*_2_AR was embedded in a 1-palmitoyl-2-oleoyl-sn-glycero-3-phosphocholine (POPC) lipid bilayer and solvated with explicit water molecules. The mutant systems were prepared by appropriately changing the residues in the wild-type receptor: Y219^5.58^A for the single mutant and T68^2.39^F, Y132^3.51^G, and Y219^5.58^A for the triple mutant.^17,18^ Sodium and chloride ions were added to neutralize and maintain the physiological concentration of 150 mM in all the systems. The CHARMM36m force-field parameters were used for the receptor,^52^ CHARMM36 for the lipid and ions,^53^ and CHARMM-modified TIP3P model was used for water.^54^ Further, the CHARMM General Force Field (CGenFF) parameters were assigned to the agonist BI-167107 using the ParamChem server,^55,56^ After energy minimization and equilibrations involving a step-by-step slow release of restraints, 100 ns conventional MD (cMD) equilibrium simulations were performed for wild-type and mutant systems. The CHARMM-GUI web server was used for system preparations,^57,58^ and AMBER18 on graphical processing units (GPUs) was used for simulations.^59^

### Gaussian-accelerated MD simulation

The conformational changes of interest in large biomolecular systems usually occur in the microsecond to millisecond time scales and are inaccessible to cMD simulation.^48,49^ To address this problem, we used GaMD, a collective variable-free enhanced sampling technique that accelerates the conformational transitions of proteins by adding a harmonic boost potential, Δ*V*(***r***), to the system potential, *V*(***r***), to overcome the energy barrier,^48,49,60^

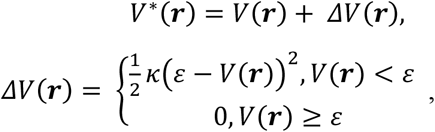

where *V*^∗^(***r***) is the modified system potential, ***r*** ={***r***_**1**_, ***r***_**2**_, **…**, _***r*N**_} is the coordinate vector of *N* atoms in the system, *κ* is the harmonic force constant, and *ε* is the threshold cut-off for *V*(***r***), below which the boost potential is added. More information about the GaMD method can be found in Refs 48 to 50. For clarification, the essential details are provided in SI methods.

The GaMD simulations were performed using the GaMD module implemented in AMBER18 software on GPUs.^50,61^ We used the standard GaMD simulation protocol described in previous studies on GPCRs.^44,62–65^ Starting from the last frame of the previous equilibration step of each system, a 12 ns short cMD was run to collect potential statistics, e.g., the maximum (*V*_*max*_), minimum (*V*_*min*_), average (*V*_*avg*_), and standard deviation (*σ*_*V*_) of system potential energies (see SI methods). Subsequently, a 40 ns GaMD equilibration was performed after adding the boost potential. We then randomized the initial atomic velocities for production simulations and started three independent trajectories of 1.1 µs for wild-type and mutant systems (a total of ~10 µs). The first 100 ns of each run was further considered as equilibration and discarded in the final analyses. A 2 fs timestep was used for GaMD, and the trajectory frames were written at every 2 ps. We used the _“_dual boost_”_ level (applying boost potential to the dihedral energy term and the total potential energy term) of GaMD after setting the reference energy to the lower bound, i.e., *ε* = *V*_*max*_. The average and standard deviation of the boost potentials were calculated at every 400,000 steps, and σ_0_ (the upper limit of standard deviation) was set to 6.0 kcal/mol for both potential and dihedral energetic terms. *PyReweighting* toolkit^65^ was used for reweighting the distributions, and the free energy plots were constructed using in-house MATLAB scripts.

### Nonbonding interaction energy estimation

The average nonbonding interaction energies of the agonist BI-167107 and the protein residues were calculated as the sum of their average electrostatic and van der Waals interaction energies over the simulation time, ⟨*E*⟩ = ⟨*E*^*elec*^ ⟩ + ⟨*E*^*vdW*^⟩. The difference in ⟨*E*⟩ between systems were calculated as Δ*E* = ⟨*E*⟩^*i*^ − ⟨*E*⟩^*j*^, where *i* and *j* denote different systems. All interaction energies were calculated using the *CPPTRAJ* module of AMBER18.^66^

### Machine learning-based clustering and classification

For conformational clustering based on root mean square deviations (RMSD) of C_α_ atoms in TM helices and H8, we utilized the Gaussian mixture model (GMM), an unsupervised machine learning (ML) algorithm that efficiently clusters complex data. The two-dimensional clustering of χ1 torsional angles of R131^3.50^ and Y326^7.53^ was performed using the *k*-means clustering algorithm. To identify critical interactions contributing to the separation of these torsional angles into distinct clusters, we utilized four machine learning classifiers, namely gradient boosting, extreme gradient boosting (XGBoost), random forest, and the automated approach AutoGluon developed by Amazon Web Services.^67^ The Xgboost^68^ Python module was used for implementing the XGBoost classifier and the Scikit-learn^69^ library for random forest and gradient boosting. For XGBoost, all parameters were set to their default values except for a learning rate of 0.3 and a maximum tree depth of 6. In the case of the random forest classifier, 25 decision trees were employed alongside the default parameters. The electrostatic as well as van der Waals pair interaction energies involving R131^3.50^ or Y326^7.53^ (with |⟨*E*⟩| ≥ 1 kcal/mol) were considered as features for all machine learning classifiers, and class labels were assigned for torsional angle clusters identified using the *k*-means algorithm. We computed the average feature importance (i.e., the importance of interaction pairs) through 10-fold cross-validation for each model and rank-ordered the residue pairs contributing up to 80% importance. The interactions of R131^3.50^ and Y326^7.53^ that emerged as important features in three out of the four models were considered as the critical residues involved in stabilizing distinct side chain conformational states for each system. Further details of ML-based methods, including performance evaluation of models, are provided in the SI text.

### Inter-residue contacts analysis

A contact between two residues is established if any two heavy atoms of the residues fall within a 4.5 Å distance for at least 75% of the simulation time. A Python script utilizing MDTraj modules^70^ was written for contact analysis and made available as a parallelizable command line program named *trajcontacts* (https://github.com/rkmlabiiserb/trajcontacts).

### Dynamic residue networks and suboptimal paths

Dynamic residue networks (DRNs) constructed from MD simulation data are extensively used to study the allosteric communications between distant residues in proteins and nucleic acids.^71–74^ Networks are constructed by considering C_α_ atoms of the receptor and the central nitrogen atom of the agonist BI-167107 (named as _‘_NAP_’_ in the active crystal structure, PDB ID: 3SN6) as nodes. The edges between nodes are defined by inter-residue contacts^71,72^ and are weighted using *d*_XY_ = −*log*(*r*_XY_), where *r*_XY_ is the generalized correlations between residue pairs X and Y.^75^ Fluctuations of atoms that represent the nodes are used to calculate generalized correlations as, 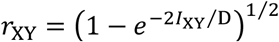, where *l*_XY_ is the mutual information between X and Y (see SI text), and D = 3 for (x, y, z) dimensions.^75^ We used *g_correlation* for calculating *r*_XY_ values.^75^

The length of a path between two distant nodes is defined as the sum of edge weights along that path (*l* = *∑*_XY_ *d*_XY_, where *d*_XY_ is the edge weight of all node pairs in the path).^76,77^ An optimal or shortest path (*l*_0_) between two nodes is the one with the least length and corresponds to the maximum correlation. Furthermore, suboptimal paths (SOP) between any two nodes are the additional paths connecting them and have lengths, *l*_SOP_ ≤ *l*_0_ + *l*_cut_, where *l*_cut_ is an arbitrary distant cut-off. Herein, we determined the number of suboptimal paths (*n*_SOP_) by choosing *l*_cut_ = 1_0_ and 15. SOPs were calculated using the *subopt* program implemented in the *NetworkView* plugin in VMD.^71,76–78^

In addition, distances and dihedral angles were calculated using the *CPPTRAJ* module of AMBER18,^66^ and volume analysis was carried out using POVME 3.0 tool.^79^ Trajectories were visualized using VMD,^78^ and images were rendered using VMD and PyMol.^80^ Plots were generated using in-house Python and MATLAB scripts and snake-plots using the online template provided by GPCRdb.^81^ More details about the system setup and the MD simulation protocol, reweighting the probability distributions of reaction coordinates, and identification of receptor residues involved in transducer contacts are provided in SI text.

## Results and Discussion

In this study, we generated three independent GaMD trajectories for each of the three *β*_2_AR systems: wild-type, single mutant Y219^5.58^A, and triple mutant T68^2.39^F, Y132^3.51^G, and Y219^5.58^A (hereafter referred to as B2RWT, B2RY, and B2RTYY, respectively), as described in Table S1 along with their average boost potentials. Unless otherwise specified, the results presented herein are based on the average over triplicate simulations. The simulated system, consisting of the receptor immersed in a POPC bilayer, solvated in explicit water, and neutralized using 150 mM NaCl, is depicted schematically in Figure S1.

### Mutations induce characteristic conformations at the extracellular and intracellular regions of the receptor

GPCRs are known to sample characteristic conformations upon selective coupling to their transducers.^12,13,16,26,33,51,82^ To understand the extracellular and intracellular variations of the transmembrane helices due to single and triple mutations, we clustered the snapshots of individual systems based on the root mean square deviations (RMSD) of their C_α_ atoms using Gaussian mixture model (GMM), an unsupervised machine learning algorithm utilized for clustering complex data (see SI methods for details). An optimal number of three clusters is formed for B2RWT and B2RTYY, and two for B2RY corresponding to minimum Bayesian information criterion scores^83^ (Figure S2). The clusters of B2RWT comprise 43.6%, 41.4%, and 15% of the population, while B2RY is clustered into 64% and 36%. The three clusters of B2RTYY have populations of 49.9%, 35.5%, and 14.6%. The larger conformational variations observed for the wild-type and triple mutant indicate greater flexibility of transmembrane regions in these systems compared to the single mutant.

In our simulation, the wild-type system shows large conformational variations at the extracellular part compared to both the active (PDB ID: 3SN6) and inactive (PDB ID: 2RH1) crystallographic structures (Figure 1a). Notably, we observe inward movements of TM2 and TM6 as well as outward movements of TM5, TM7, and extracellular loop 2 (ECL2) with respect to the helical core in the first cluster of B2RWT. For the single mutant, TM6 shifts inward while TM7 moves outward (Figure 1b). The triple mutant also exhibits an inward movement of TM6 and a shift of ECL2 towards TM5 (Figure 1c). We note that the TM6 inward movement commonly observed across all three systems represents a conformation observed in arrestin-bound structures of the avian *β*_1_AR (56% sequence identity to *β*_2_AR),^82^ and rhodopsin.^84^ Other clusters of all three systems mostly display extracellular conformations comparable to their primary cluster, as shown in Figure S3.

**Figure 1.**
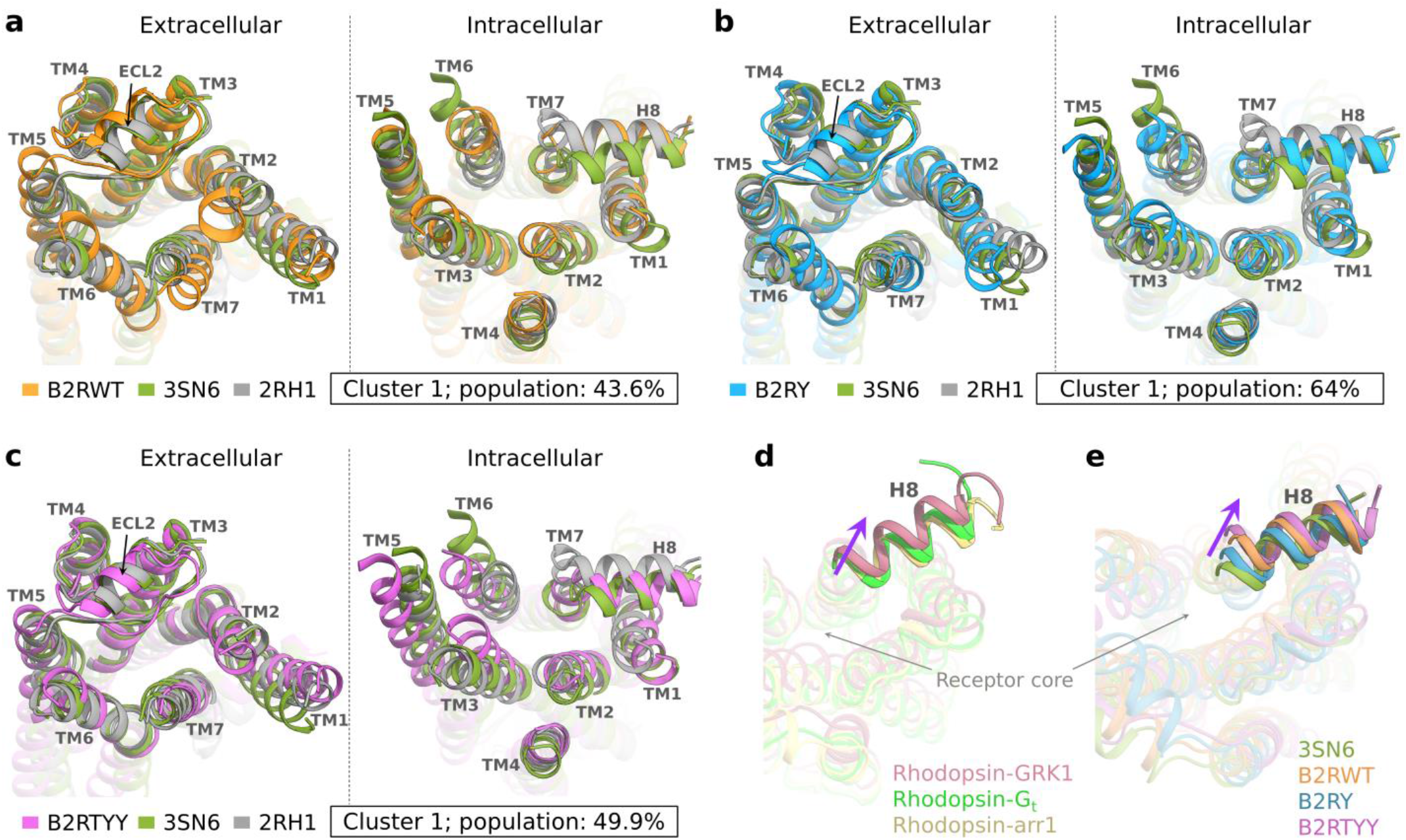
**(a-c)** Conformational variations in the extracellular and intracellular regions of *β*_2_AR due to mutations. Representative snapshots extracted from clusters of each system are superimposed with the active state starting structure (PDB ID: 3SN6) and an inactive state structure (PDB ID: 2RH1). Cluster numbers, along with their populations, are given inside boxes. Intracellular view of H8 conformations in **(d)** rhodopsin bound to different transducers (rhodopsin-GRK1, PDB ID:7MTA; rhodopsin-G_t_, PDB ID:6OYA; and rhodopsin-arr1, PDB ID:5W0P), and **(e)** structures representing peaks in the distribution of H8 displacement from the receptor core (shown in Figure S7e) for all three systems. Thick violet arrows denote the outward shift of H8 in rhodopsin-GRK1, B2RWT, and B2RTYY.

At the cytosolic part, the outward shift of TM6 from the receptor core is one of the known structural hallmarks of GPCR activation.^28,51^ Our simulations of the transducer-free wild-type receptor starting from the active crystal structure exhibit intermediate TM6 conformations (Figures 1a, S3a, and S3b). The cluster-wise and overall distributions of TM6 displacement from the inactive structure (measured at E268^6.30^ C_α_) for the wild-type and mutant systems are provided in Figure S4. It is to be noted that the fully active state of the receptor with a larger outward shift of intracellular TM6 can only be attained in the presence of the bound transducer, as observed in previous NMR and MD simulation studies of *β*_2_AR.^85,86^ Additionally, TM3 moves inward, and TM2 positions identically to that in the inactive structure for wild-type, indicating the collapse of the intracellular transducer-binding cavity compared to the active structure. Consequently, the average volume of this cavity decreases to 348 ± 16 Å^3^ in the wild-type from 118^3^ Å^3^ in the active structure (Figure S7a). Note that the intracellular cavity volume of the inactive receptor is measured to be 269 Å^3^.

For the single mutant, intracellular TM6 shows a relatively smaller inward shift from the starting active structure compared to the wild-type (Figures 1b, S3c, and S4); moreover, the outward shift of TM5 from the helical core (measured at L230^5.69^ C_α_ as explained in Figure S5), and the position of TM2 close to the active rather than inactive structure result in a higher volume (754 ± 9 Å^3^) of the transducer-binding cavity (Figure S7b) compared to the wild-type. For the triple mutant, TM6 exhibits a similar shift as in the single mutant; however, its position is closer to TM5, and is even more pronounced in the second and third clusters (Figures 1c, S3d, and S3e). We also observe a significant outward shift of TM5 from the helical core in B2RTYY (Figure S5). Furthermore, a greater inward movement of TM3 places it close to TM6 in B2RTYY (Figures 1c, S3d, S3e, and S7d). These conformational rearrangements render TM5 unavailable for lining the transducer-binding cavity, causing a reduction of its volume to 412 ± 4 Å^3^ (Figure S7c), suggesting an unlikely engagement of G proteins that require a broader intracellular cavity, compared to *β*-arrestins that bind to a narrower cleft.^82,87–90^ This further suggests that the conformational alteration of TM5 observed in the triple mutant is unfavorable for establishing interactions between the α5 helix of G_s_ and TM5 as in the *β*_2_AR-G_s_ crystal structure.^51^ Notably, a similar TM5 conformation was also observed in our previous MD simulation study of a phosphorylated *β*_2_AR that sampled *β*-arrestin-favoring conformations over G_s_.^44^ Moreover, we examined the conformational variations of the conserved NPxxY motif at the intracellular TM7 for the wild-type and mutant systems. This motif is reported to rearrange its conformation due to a partial unwinding of TM7 around Y326^7.53^ and an inward movement of the helix towards the receptor core.^91,92^ The backbone RMSDs of the NPxxY motif are found to be similar among all three systems, with differences in peak values measuring within 0.2 Å (Figure S6).

Furthermore, we also investigated conformational changes in helix 8 (H8), as this intracellular helix was reported to shift away from the receptor core in the structure of the rhodopsin-GRK1 complex compared to its other transducer complexes (Figure 1d).^93^ B2RY that inhibits GRK binding shows a minimum deviation of H8 from the receptor core of the active crystal structure, while B2RWT and B2RTYY exhibit large displacements (Figures 1e and S7e). This suggests that the specific positioning of H8 may play a vital role in the GRK engagement of *β*_2_AR, like that in rhodopsin. In fact, a previous study based on chemical cross-linking and mass spectrometry also indicated the involvement of H8 residues in *β*_2_AR-GRK5 interactions.^94^

In addition to the TM helices and H8, intracellular loop 3 (ICL3) that connects TM5 and TM6 is reportedly involved in transducer binding to GPCRs and biased signaling.^51,82,88,93,95–103^ A recent study on *β*_2_AR using experiments and MD simulations has demonstrated that ICL3 acts as an autoregulator of GPCR signaling by sampling conformations, which are in a dynamic equilibrium between states that block and expose the transducer-binding cavity.^104^ To investigate the relative position of ICL3 with respect to the intracellular cavity, we measured the distance between ICL3 and H8 (L258^ICL3^ C_α_ - L340^8.58^ C_α_ distance) in different clusters of the wild-type and mutant systems (Figure 2). The wild-type clusters exhibit an ICL3-H8 median distance of ~30 Å (Figure 2: orange color), and representative structures display occlusion of the transducer-binding cavity, i.e., closed state (cluster 1: ICL3-H8 distance 29.5 Å; cluster 2: ICL3-H8 distance 30 Å, and cluster 3: ICL3-H8 distance 29 Å). A similar conformation was also reported in the balanced ligand (norepinephrine)-bound simulations of *β*_2_AR.^33^ In contrast, the primary cluster of the single mutant shows a large ICL3-H8 distance of 41 Å with an ICL3 orientation away from the intracellular cavity (i.e., open state; Figure 2: blue color), which could facilitate the G protein engagement, as has been reported in previous studies.^33,104^ It is to be noted that we observe an additional partially open state in the single mutant, characterized by ICL3-H8 distances measuring 37.7 Å (Figure 2). Whereas in the triple mutant, a large variation in ICL3-H8 distances is observed across the clusters (Figure 2: magenta color). The representative structures display closed and partially open states of the transducer-binding cavity (cluster 1: ICL3-H8 distance 32.4 Å; cluster 2: ICL3-H8 distance 27.2 Å, and cluster 3: ICL3-H8 distance 36 Å). A greater population of closed conformations potentially leads to the occlusion of the intracellular cavity for G protein binding. However, the diversity in the ICL3 position in the triple mutant indicates a complex role of the loop in transducer coupling. Notably, previous all-atom simulations of *β*_2_AR bound to *β*-arrestin-biased ligands in the absence of transducers showed closed ICL3 conformations,^33^ while coarse-grained simulations of the *β*_2_AR-*β*-arrestin 2 complex exhibited interactions between ICL3 and *β*-arrestin 2 under acidic membrane conditions.^105^ Moreover, the structure of *β*-arrestin 1 complexed with neurotensin receptor 1 also revealed the involvement of ICL3 in the arrestin-receptor interface.^96^ Thus, considering previous findings, the increased dynamics of ICL3 in the *β*-arrestin-favoring triple mutant, compared to other systems, likely represents a specific signature of *β*-arrestin binding. Taken together, a comparison of the conformational ensembles of the mutant receptors with the wild-type reveals that distinct rearrangements of transmembrane helices, particularly TM3, TM5, TM6, and TM7, along with H8 and ICL3 at the cytosolic side, could determine the experimentally observed transducer selectivity.

**Figure 2.**
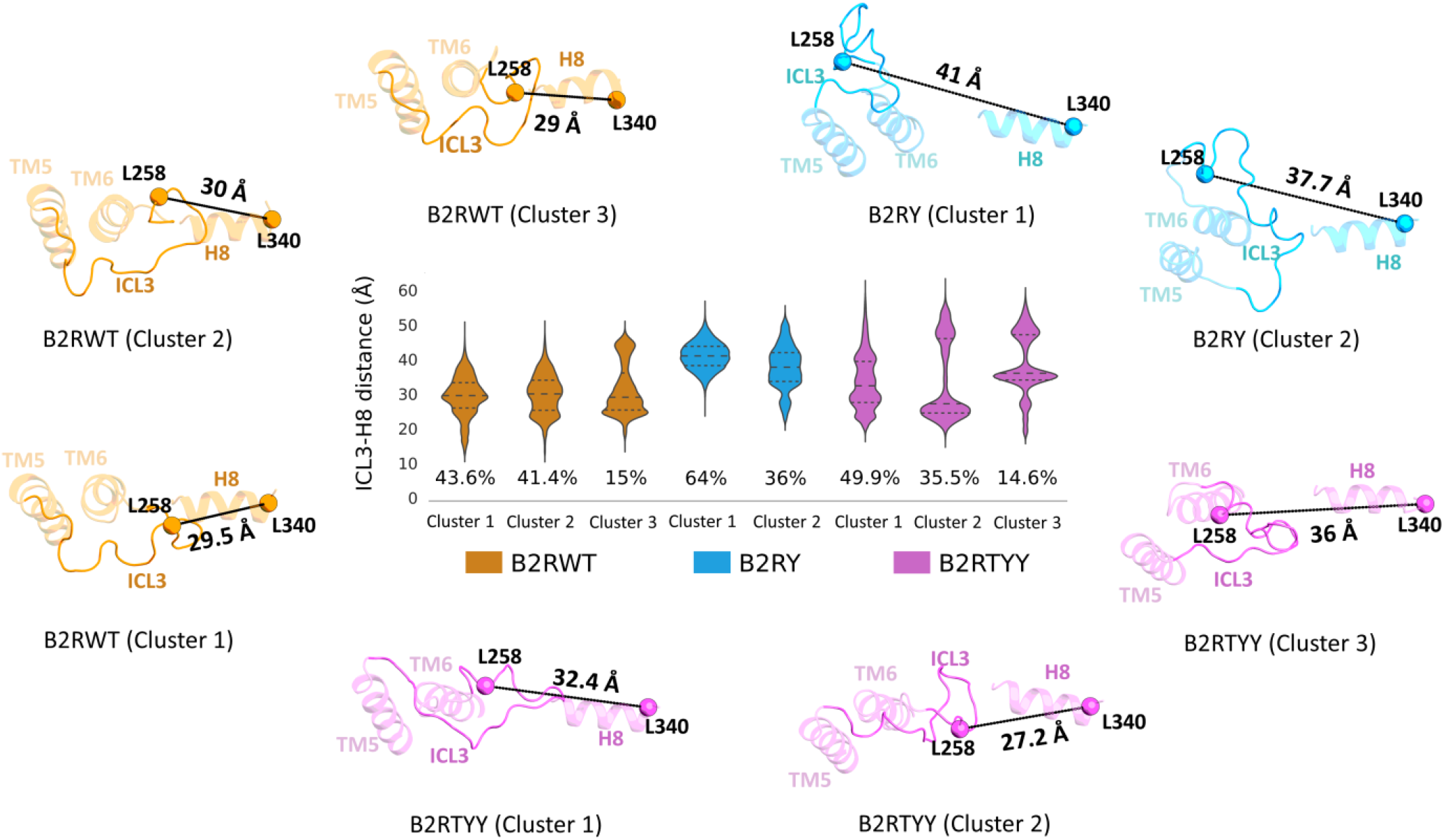
Violin plots (center) depicting the cluster-wise distributions of ICL3-H8 distance (L258^ICL3^ C_α_ - L340^8.58^ C_α_). Violins are marked with quartiles using discontinuous lines (median at the middle), and the population percentage in each cluster is given at the bottom. Representative snapshots having median values of ICL3-H8 distance for all clusters are provided in the same colors as the violins

### Mutant receptors favor specific rotameric states of the conserved intracellular residues R131^3.50^ and Y326^7.53^

Fine-tuning the conformations of conserved residues located at various regions of a GPCR is crucial in the modulation of transducer-specific signaling.^13,24,30^ Recent studies showed that at the intracellular part, the arginine residue R^3.50^ of the D(E)RY motif in TM3 and the tyrosine residue Y^7.53^ of the NPxxY motif in TM7 (Figure S8a) have rearranged their conformations to sample transducer selective states of GPCRs.^26,34^ In particular, MD simulations of angiotensin II type 1 receptor (AT1R) revealed sampling of downward rotameric states of R^3.50^ and Y^7.53^ (side chains point towards the cytosolic region), corresponding to an alternative active state that prefers *β*-arrestin binding as opposed to the upward rotamers favoring G protein.^26^ Moreover, the modeled structures of transducer-bound AT1R, together with the simulations of arrestin-1-bound rhodopsin, suggested that the downward rotameric states of these conserved residues favorably interact with arrestin while R^3.50^ sterically clash with G protein.^26^ Consistently, in G_s_-bound structures of *β*_2_AR^51^ and closely related *β*_1_AR,^106^ both the side chains of R^3.50^ and Y^7.53^ adopt upward rotamers as shown in Figures S8b and S8c. Due to the unavailability of the *β*-arrestin-bound *β*_2_AR structure, we examined *β*_1_AR and observed that the finger loop of *β*-arrestin-1 favorably interacts with the downward rotamer of R^3.50^ (Figure S8c),^82^ suggesting the possibility of similar interaction for *β*_2_AR. In addition, the *β*-arrestin 1-bound structure of serotonin 5-HT_2B_ receptor^95^ displays an upward rotamer of the side chain of R^3.50^, indicating this rotameric state of class A GPCRs may as well be compatible to form a stable complex with *β*-arrestins (Figure S8e).

We investigated the side chain conformations of R131^3.50^ and Y326^7.53^ for the wild-type and mutant *β*_2_AR systems and found two conformational states, viz., upward (↑) and downward (↓) rotamers of these residues (Figure 3). The two-dimensional (2D) potential of mean force (PMF) as a function of χ1 torsional angles of R131^3.50^ and Y326^7.53^ displays four minima in each system, as shown in Figures 3a-3c. The R↓ and R↑ conformers peak around 180º and 295º, respectively, while Y↓ and Y↑ are close to 175º and 290º, respectively (Figures 3a-3c; side and top panels). To identify the difference in populations of the observed conformational states, we performed a 2D *k*-means clustering^107^ over the entire ensemble for each system (see SI methods for details). The wild-type system is found to form an optimal number of three clusters, while the mutant systems exhibit four clusters, corresponding to the maximum silhouette coefficient scores^108^ (Figure S8), and the populations of the obtained clusters are given in Table 1. It is observed that R131^3.50^ predominantly samples upward rotamers in all systems. The states R↑–Y↓ and R↑–Y↑ together constitute the majority of the conformations, with a combined population of 97.3% in B2RWT, 89.2% in B2RY, and 76.2% in B2RTYY. The larger conformational populations of R↑ in the wild-type and single mutant suggest that these systems can favorably interact with G protein compared to the triple mutant. Moreover, relatively smaller energy barriers between states R↑–Y↓ and R↑–Y↑ in the wild-type and single mutant indicate that G protein could bind with both the rotameric states of Y326^7.53^ of *β*_2_AR (Figures 3a and 3b).

**Table 1.** Populations of rotameric states of R131^3.50^ and Y326^7.53^ clustered using the *k*-means method. Four major clusters are identified in the mutated systems B2RY and B2RTYY, namely, R↓–Y↓, R↓–Y↑, R↑–Y↓, and R↑–Y↑. B2RWT forms three clusters, with the conformational population of the R↓–Y↓ and R↓–Y↑ states constituting a single group.

| Cluster name | B2RWT |  | B2RY |  | B2RTYY |  |
| --- | --- | --- | --- | --- | --- | --- |
|  | Cluster centroid | Cluster population (%) | Cluster centroid | Cluster population (%) | Cluster centroid | Cluster population (%) |
| R↑-Y↓ | (291.3, 184.9) | 73.2 | (290.7, 187.6) | 27.8 | (290.6, 178.1) | 17.9 |
| R↑-Y↑ | (292.5, 286.0) | 24.1 | (292.1, 291.3) | 61.4 | (291.3, 297.9) | 58.3 |
| R↓-Y↓ | (185.2, 222.3) | 2.7 | (181.0, 192.2) | 2.1 | (185.3, 176.6) | 4.5 |
| R↓-Y↑ | | | (182.3, 290.6) | 8.7 | (184.4, 295.2) | 19.3 |

**Figure 3.**
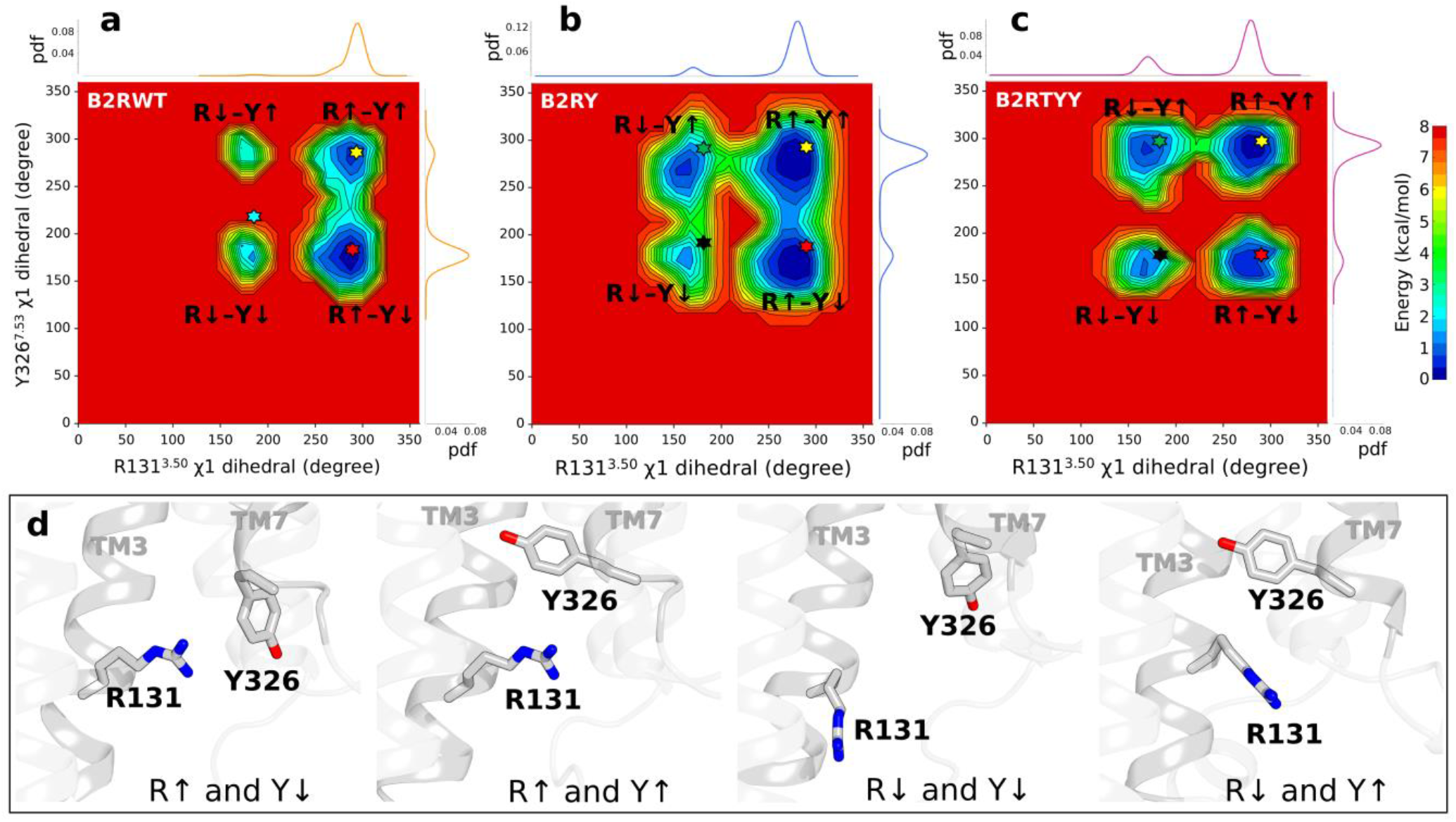
**(a-c)** Two-dimensional PMF profiles as a function of χ1 torsions of R131^3.50^ and Y326^7.53^ for all systems. The distributions of individual collective variables are shown as side and top panels of each free energy plot. Stars denote the central structures obtained from *k*-means clustering. **(d)** Representative snapshots displaying all four states observed in the PMF calculations.

The analysis of the downward states of R131^3.50^ reveals that the triple mutant exhibits the highest prevalence of R↓–Y↓ state, with a population more than two times that of the wild-type and single mutant (Table 1). This suggests a coupled downward rotation of χ1 torsions of R131^3.50^ and Y326^7.53^ is a *β*-arrestin-favoring structural phenotype of *β*_2_AR similar to that in AT1R.^26^ However, the high abundance of the R↓–Y↑ state in B2RTYY indicates a possible alternative active state of *β*_2_AR that prefers *β*-arrestin over G protein with an upward rotamer of Y326^7.53^ as has been observed for κ-opioid receptor recently.^34^

To understand the energetic influence of mutated residues (T68^2.39^F, Y132^3.51^G, and Y219^5.58^A) on the rotameric residues, we estimate their average nonbonding interaction energies, ⟨*E*⟩. The tyrosine residue Y219^5.58^ in B2RWT exhibits favorable interactions with both R131^3.50^ and Y326^7.53^ (Figure S10a). The interaction with R131^3.50^ is mainly long-range electrostatic, whereas its distance with Y326^7.53^ indicates the formation of water-mediated hydrogen bonds in a major fraction of simulations (Figure S10b), as was observed in the active state of *β*_2_AR.^27,51,109^

These connections are abolished due to Y219^5.58^A mutation in both B2RY and B2RTYY (Figure S10a), leading to an increase in flexibility of their sidechains, thereby potentially allowing to sample a broader conformational space. In B2RTYY, the introduction of a bulky sidechain via the T68^2.39^F mutation increases the van der Waals interactions with R131^3.50^ and Y326^7.53^ (Figure S10a), possibly contributing to a greater prevalence of downward rotameric states compared to other systems.

Next, to discern the molecular basis of characteristic conformational states involving the downward rotamer of R131^3.50^ (in R↓–Y↓ and R↓–Y↑), hereafter referred as R↓ states, which are likely to be important in transducer (*β*-arrestin) specificity,^26,34,82^ we further examine the critical interactions formed by the rotameric residues with other receptor residues. For identifying the specific interactions that distinguish different states, we employ four machine learning classification algorithms — gradient boosting, XGBoost, random forest, and AutoGluon. The nonbonding energies (electrostatic as well as van der Waals) of interactions involving R131^3.50^ or Y326^7.53^ for all simulated frames are considered as features, and their cluster affiliations as class labels. Top residue pairs that effectively predict the clusters for the test (unseen) data in each system are extracted, as detailed in the Materials and Methods section. Interactions that stabilize R↓ or R↑ states are shown in Figure 4 and their numerical values are provided in Table 2.

**Table 2.** Numerical values of average nonbonding energies for interactions of R131^3.50^ with the mutated residues, and critical residues in upward (R↑) and downward (R↓) rotameric states identified using machine learning classifiers.

| B2RWT |  |  | B2RY |  |  | B2RTYY |  |  |
| --- | --- | --- | --- | --- | --- | --- | --- | --- |
| Residue pairs | $\langle E \rangle$ (kcal/mol) | | Residue pairs | $\langle E \rangle$ (kcal/mol) | | Residue pairs | $\langle E \rangle$ (kcal/mol) | |
| | $R_{\uparrow}$ | $R_{\downarrow}$ | | $R_{\uparrow}$ | $R_{\downarrow}$ | | $R_{\uparrow}$ | $R_{\downarrow}$ |
| $R131^{3.50}$-T68<sup>2.39</sup> | -1.5 | 0.03 | $R131^{3.50}$-T68<sup>2.39</sup> | -1.45 | 0.02 | $R131^{3.50}$-F68<sup>2.39</sup> | -1.82 | -1.73 |
| $R131^{3.50}$-S120<sup>3.39</sup> | -1.25 | -1.14 | $R131^{3.50}$-S120<sup>3.39</sup> | -1.53 | -1.47 | $R131^{3.50}$-T123<sup>3.42</sup> | -1.31 | -1.05 |
| $R131^{3.50}$-T123<sup>3.42</sup> | -1.37 | -1.13 | $R131^{3.50}$-I121<sup>3.40</sup> | -1.06 | -1.01 | $R131^{3.50}$-L124<sup>3.43</sup> | -2.07 | -1.85 |
| $R131^{3.50}$-I127<sup>3.46</sup> | -8.68 | -7.75 | $R131^{3.50}$-T123<sup>3.42</sup> | -1.2 | -1.1 | $R131^{3.50}$-I127<sup>3.46</sup> | -8.55 | -7.71 |
| $R131^{3.50}$-I135<sup>3.54</sup> | -2.97 | -2.9 | $R131^{3.50}$-L124<sup>3.43</sup> | -2.18 | -2.05 | $R131^{3.50}$-I135<sup>3.54</sup> | -2.7 | -2.9 |
| $R131^{3.50}$-Y219<sup>5.58</sup> | -1.99 | -2.16 | $R131^{3.50}$-I127<sup>3.46</sup> | -8.35 | -7.67 | $R131^{3.50}$-A219<sup>5.58</sup> | -0.09 | -0.01 |
| $R131^{3.50}$-D251<sup>ICL3</sup> | -21.63 | -21.73 | $R131^{3.50}$-Y141<sup>ICL2</sup> | -1.29 | -0.48 | $R131^{3.50}$-E268<sup>6.30</sup> | -20.93 | -21.92 |
| $R131^{3.50}$-I325<sup>7.52</sup> | -0.67 | -0.94 | $R131^{3.50}$-A219<sup>5.58</sup> | -0.29 | -0.42 | $R131^{3.50}$-A271<sup>6.33</sup> | -1.5 | -1.74 |
| | | | $R131^{3.50}$-E249<sup>ICL3</sup> | -31.78 | -27.78 | $R131^{3.50}$-L275<sup>6.37</sup> | -0.75 | -0.58 |
| | | | $R131^{3.50}$-D251<sup>ICL3</sup> | -24.22 | -27.97 | | | |
| | | | $R131^{3.50}$-T274<sup>6.36</sup> | -7.1 | -5.62 | | | |

**Figure 4.**
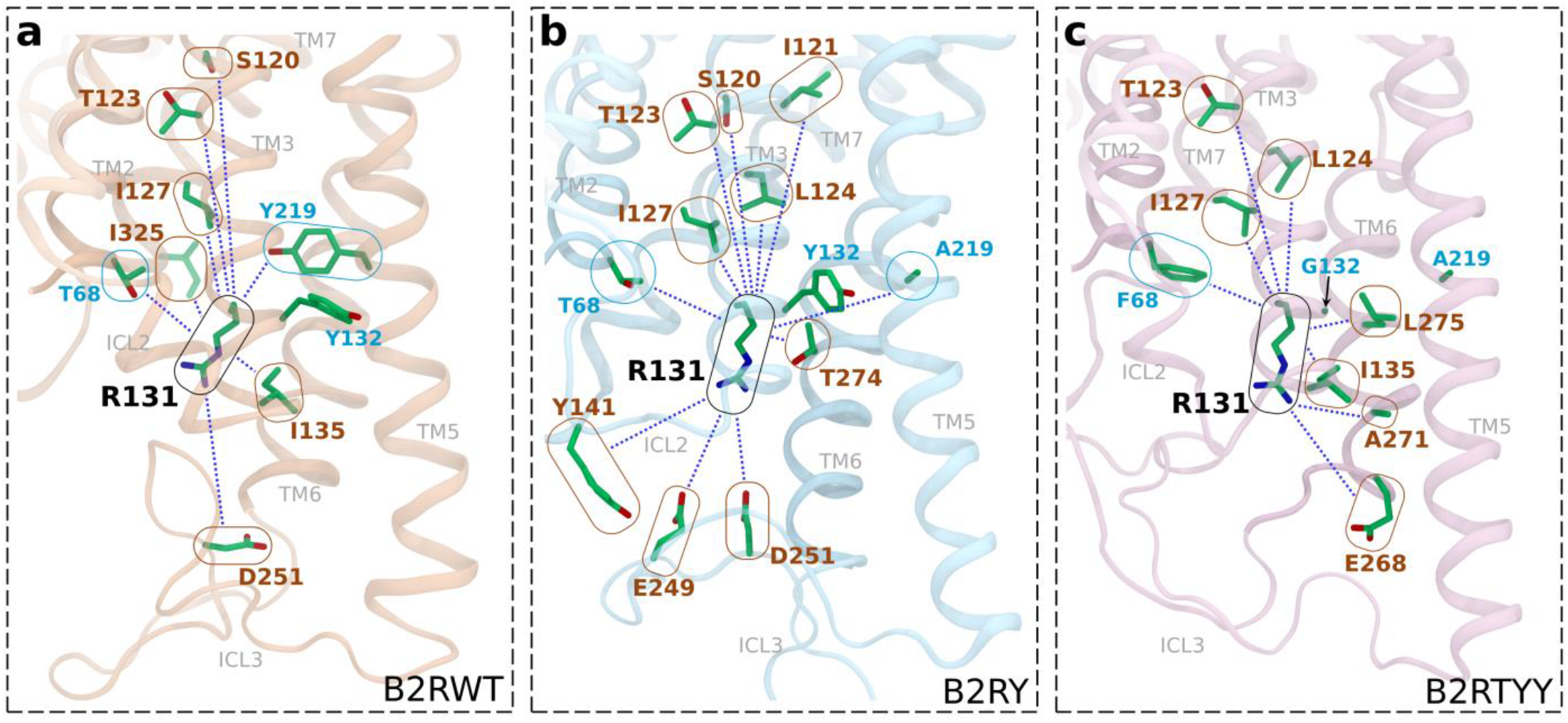
Representative snapshots (of the R↓ states) showing the favorable nonbonding interactions of R131^3.50^ with mutated residues and other critical residues that significantly contribute to the formation of distinct clusters for **(a)** B2RWT, **(b)** B2RY, and **(c)** B2RTYY. Blue dotted lines represent interactions, and residues are marked using circles/ovals: R131^3.50^ in black, mutational site residues in cyan, and other critical residues in brown. The numerical values of the average interaction energies in upward and downward rotameric states are provided in Table 2.

The inter-residue interactions involving R131^3.50^ primarily stabilize its upward rotamers, which constitute the largest conformational population in each system (Tables 1 and 2). However, the transient formation of R↓ states in B2RWT (only 2.7% of the population) is facilitated by the presence of favorable interactions at the cytosolic end, including that with ICL3 residue D251^ICL3^ and TM3 residue I135^3.54^ (Figure 4a). Notably, the favorable interaction with TM2 residue T68^2.39^ that stabilize the upward rotameric states of R131^3.50^ is abolished. Additionally, the stabilizing interactions with TM3 residues I127^3.46^, T123^3.42^, and S120^3.39^, located above R131^3.50^ (towards the extracellular end with respect to the spatial location of R131^3.50^), are marginally decreased in the R↓ states (Table 2). Rearrangements of this energy network upon single and triple mutations are likely to contribute to the increased populations of R↓ states in B2RY and B2RTYY.

As discussed, the loss of critical interaction of R131^3.50^ with Y219^5.58^A in B2RY drives larger conformational sampling of R↓ states (10.8%) as compared to B2RWT. In fact, interactions with several TM3 residues (I127^3.46^, L124^3.43^, T123^3.42^, I121^3.40^, and S120^3.39^) and TM6 residue T274^6.36^ are weakened, thereby promoting R↓ states (Figure 4b and Table 2). Moreover, on the cytosolic side, we observe increased stabilizing interactions of R131^3.50^ with ICL3 residue D251^ICL3^ (Table 2). The significant increase in downward rotamers of R131^3.50^ (23.8%) in B2RTYY can largely be attributed to its favorable interaction with the mutated residue F68^2.39^, which was absent in B2RWT (Table 2). Additional stabilizing interactions with E268^6.38^, A271^6.33^, and I135^3.54^ on the cytosolic side and reduced interaction strength of I127^3.46^, L124^3.43^, and T123^3.42^ towards extracellular side contributed to the overall increase in R↓ population. Notably, E268^6.38^ is a highly conserved residue that reproducibly forms salt-bridge interactions with R131^3.50^ in the inactive state of *β*_2_AR.^110^ However, the salt-bridge interaction has not been observed in any of the systems considered in this study (Figures S10b-S10d), suggesting that our simulated conformational ensembles do not resemble the inactive state of *β*_2_AR. Moreover, the involvement of E268^6.38^ in stabilizing R↓ states suggest the intricate interplay of conserved structural domains within GPCRs in conformational transitions amenable to biased signaling.

### Mutations distinctly alter the intrinsic allosteric communications between the agonist and intracellular region of the receptor

The existence of allosteric communications between the intracellular and extracellular regions is well-known for GPCRs.^111–114^ The strengths of these communications are correlated with the binding affinities of both orthosteric ligand and intracellular transducers.^28,113,115^ The conformational variations observed at the extracellular and intracellular receptor regions upon mutations suggest that the underlying intrinsic allosteric communications primed by agonist binding are altered. To investigate the impact of transducer-selective mutations on interactions of the agonist BI-167107 with the receptor residues, their average nonbonding energies, ⟨*E*⟩, are computed, and the corresponding differences in the energies (Δ*E*) between the mutated and wild-type systems are shown in Figure S11. We observe alterations in agonist interaction energy for several orthosteric pocket residues in B2RY and B2RTYY (Figures S11a and S11b), and it clearly establishes the allosteric perturbations at the distal extracellular regions induced by transducer selective mutations. We note that previous experimental and MD simulation studies have highlighted the role of extracellular agonist-binding residues in modulating the biased signaling in GPCRs.^116–119^

To understand the system-specific variations in agonist interactions due to mutations, we calculated Δ*E* between the triple mutant and single mutant (Figure 5). Residues having positive (or negative) Δ*E* values indicate the interactions with the agonist are more favorable in B2RY (or B2RTYY). Among the residues in TM3 (D113^3.32^, V114^3.33^, and V117^3.36^), TM5 (S204^5.43^ and S207^5.46^), TM6 (F290^6.52^ and N293^6.55^), and TM7 (N312^7.39^) that exhibit strong favorable interactions with the agonist in B2RY (Figure 5), several are found in proximity to the base of the orthosteric pocket. Interestingly, residues in this region are reported to play a crucial role in the G protein signaling bias of GPCRs; for instance, S204^5.43^ and N293^6.55^ are essential in the G protein-selective signaling induced by the partial agonist salmeterol for *β*_2_AR.^24,45,120^ However, it should be noted that the G protein bias of B2RY is caused by the inhibition of GRK-binding that abolishes *β*-arrestin signaling.^18^ Therefore, more favorable agonist interactions at the base of the orthosteric pocket in B2RY suggest a possible enhancement of G protein bias coupled with GRK inhibition. Whereas for the *β*-arrestin-biased B2RTYY, the residues in ECL2 (Y174^ECL2^, D192^ECL2^, F193^ECL2^, and T195^ECL2^), TM5 (Y199^5.38^), and TM7 (I309^7.36^ and Y316^7.43^), close to the extracellular space distal to the base of the orthosteric pocket, exhibit more favorable agonist interactions. Notably, the ECL2 residue F193^ECL2^ has been previously identified as crucial in *β*-arrestin signaling, as its mutation to alanine leads to a bias of *β*_2_AR away from *β*-arrestin interactions.^46,121^ In addition, the TM5 residue Y199^5.38^ has been shown to act as a microswitch that regulates the engagement of *β*-arrestin with *β*_2_AR.^19^ Moreover, structural studies on the highly homologous *β*_1_AR revealed that *β*-arrestin-biased ligands, such as bucindolol and carvedilol, establish more contacts with the extracellular residues than with the deep orthosteric pocket residues.^24,82,122^ In effect, the specific interactions of the agonist-binding residues with BI-167107 upon mutations, along with the observed conformational changes in intracellular parts, indicate differential communications between the agonist and transducer binding regions in the single and triple mutants, which are detailed further in this section.

**Figure 5.**
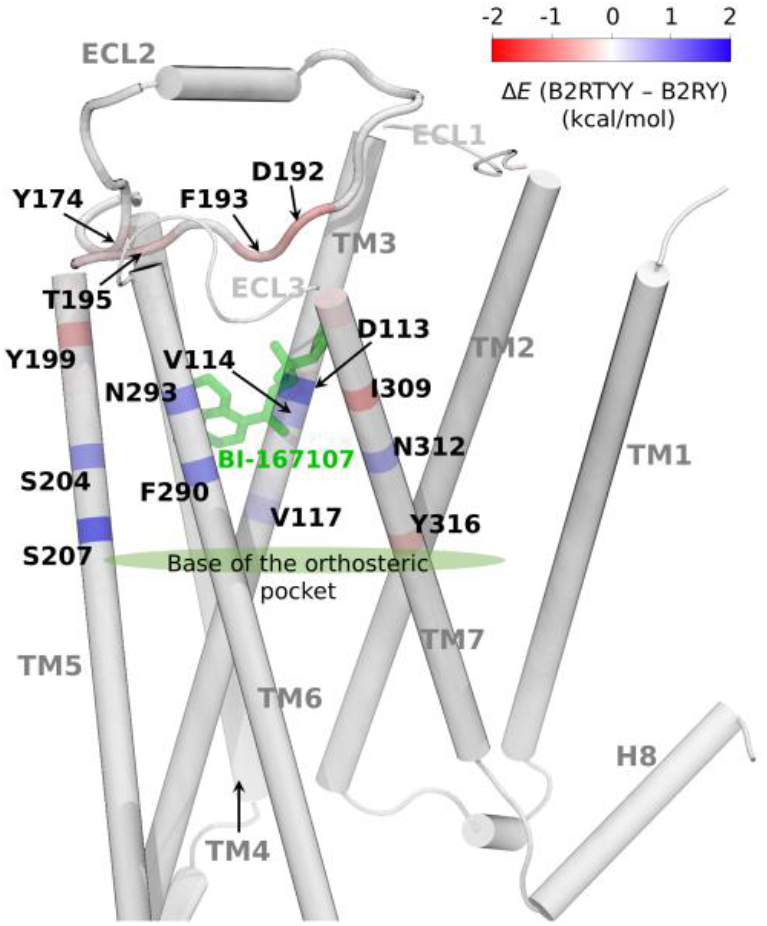
The difference in nonbonding interaction energies of the agonist BI-167107 with the receptor residues between B2RTYY and B2RY. The critical residues that engage in relatively high favorable interactions in the single mutant (blue) and triple mutant (red) are marked.

To examine the change in allosteric signal transfer between the agonist and intracellular region upon mutations, we constructed dynamic residue networks (DRNs) by considering each residue and the agonist as nodes and the contacts between the nodes as edges. The edge weights are determined using generalized correlations, and the shortest/optimal paths are calculated between the nodes. While the shortest path is typically regarded as the most biologically significant route for allosteric information transfer between a pair of nodes, it has been shown that an ensemble of paths with similar lengths, albeit slightly longer than the shortest path, can also contribute to allostery.^123^ These degenerate paths are called suboptimal paths (SOPs) and are identified using a distance cut-off *l*_cut_^76,123^ (see Materials and Methods). The number of suboptimal paths (*n*_SOP_) between two residues measures the strength of allosteric communication between them.^77^ By determining all SOPs that connect the extracellular agonist with the residues in the intracellular region, we aimed to identify the changes in allosteric communications leading to selective signaling that disrupted GRK interaction for B2RY and G_s_interaction for B2RTYY. Residues in the intracellular loops, H8, and cytosolic halves of TM1 to TM7, which are at least 10 Å apart from the agonist, were considered for the analysis. We used *l*_cut_values of 10 and 15 to compute *n*_SOP_ and found that both show a similar pattern for each system (Figures 6a and S12, and Table S3). Additionally, the identification of residues at the interface of *β*_2_AR and transducers (G protein/*β*-arrestin/GRK) is performed using a Python script – *trajcontacts* (listed in Table S4); the residues at the G protein interface are obtained from the *β*_2_AR-G_s_complex structure (PDB ID: 3SN6) and the residues at the GRK/*β*-arrestin interface are determined by aligning the crystal structure of the receptor (from 3SN6) with other available *β*-arrestin/GRK complex structuresof class A GPCRs (see SI methods for details).

**Figure 6.**
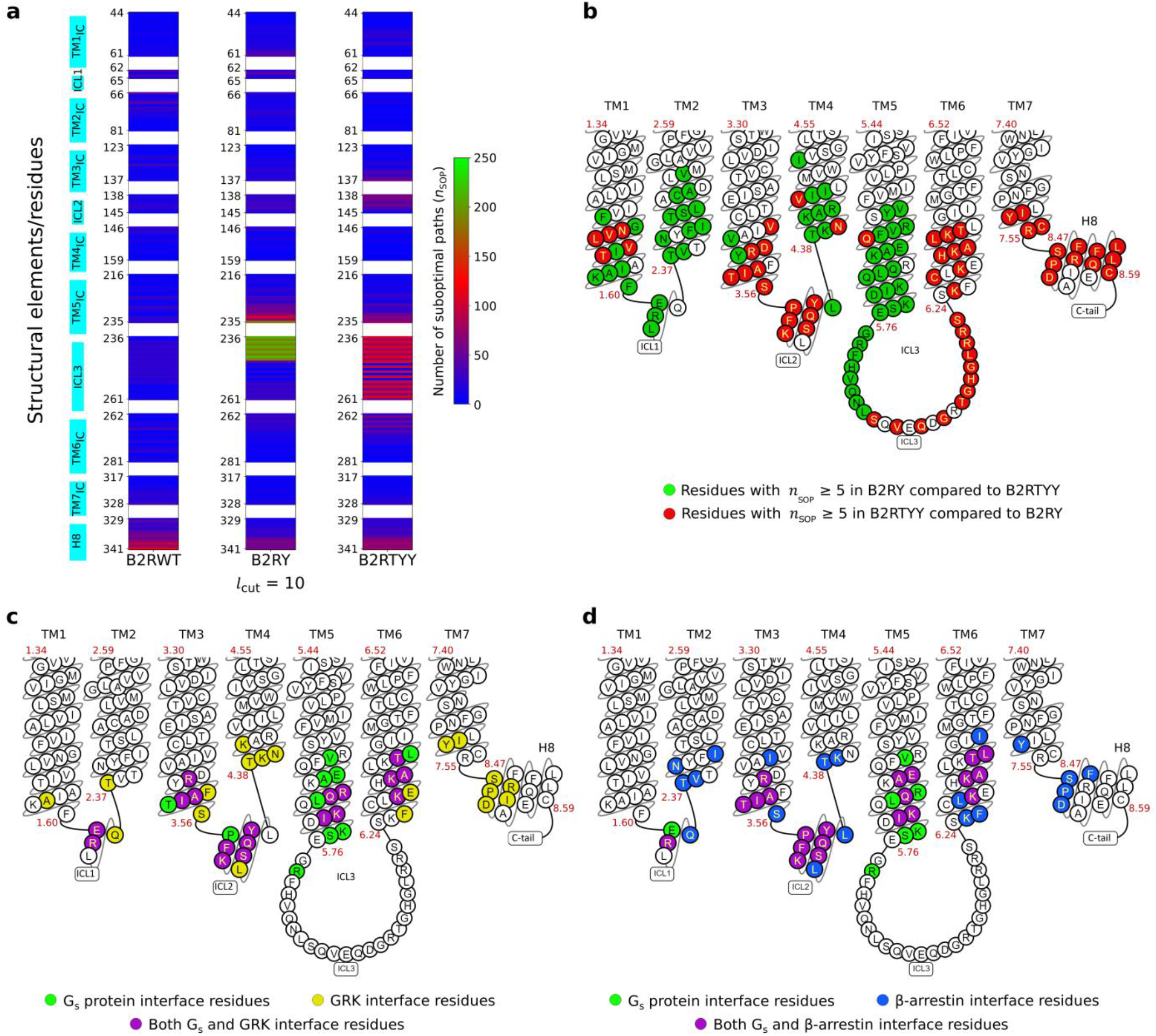
**(a)** Heatmap showing the number of suboptimal paths (*n*_SOP_), from the agonist BI-167107 to the intracellular (IC) residues (residues at the cytosolic half of the receptor that are at least 10 Å apart from the agonist) for *l*_cut_= 10. The numerical values of *n*_SOP_ are provided in Table S3. **(b)** Snake plot showing intracellular residues with *n*_SOP_ ≥ 5 for a given mutant over the other: B2RY in green, and B2RTYY in red. The residues at the interface of *β*_2_AR with **(c)** G_s_and GRK as well as **(d)** G_s_and *β*-arrestin, are marked in snake plots. The residues at the G protein interface are obtained from the *β*_2_AR-G_s_complex structure (PDB ID: 3SN6), and the residues at the *β*-arrestin/GRK interface are determined by aligning the crystal structure of the receptor (from the *β*_2_AR-G_s_complex, PDB ID: 3SN6) with other *β*-arrestin/GRK complex structures of class A GPCRs; the *β*_2_AR residues within 4 Å proximity of *β*-arrestin/GRK are identified as interface residues (see SI methods for details).

Compared to B2RWT, B2RY and B2RTYY exhibit an increase in the number of suboptimal paths from the extracellular agonist to the intracellular residues (Figures 6a, S12, and Table S3). Nevertheless, a large *n*_SOP_ is observed for several intracellular residues, especially in H8, implicating a strong allosteric coupling of these regions with the agonist in B2RWT (Figure 6a and Table S3). In B2RY, the ICLs, intracellular ends of TM1 and TM5, and H8 exhibit noticeable allosteric communications from the agonist, whereas in B2RTYY, ICL2 and ICL3, intracellular ends of several TMs (in particular, TM3, TM5, and TM6), and H8 show a substantial number of suboptimal paths (Figure 6a and Table S3). The increase in allosteric communications in the mutant receptors clearly suggests constrained intracellular dynamics, where distinctive conformational states are sampled that could favor preferential engagement of transducers.

To investigate the differences in allosteric communication from the agonist to the transducer-binding interfaces in B2RY and B2RTYY, we examined the intracellular residues with *n*_SOP_ ≥ 5 for a given mutant over the other (Figure 6b). Notably, the intracellular loops and intracellular ends of several TMs (TM1 to TM5) participate in the allosteric signal transfer in B2RY. A number of residues within these structural elements, especially in ICL1 and TM5, are found to interact with G_s_(Figure 6c and Table S4) and/or reported to be crucial in G protein signaling.^51,124,125^ In B2RTYY, the ICL2, ICL3, H8, and the intracellular ends of many TMs (particularly TM3, TM6, and TM7), are allosterically coupled with the agonist (Figure 6b), where several residues have been identified to be part of the modeled GRK/*β*-arrestin interface of *β*_2_AR (Figures 6c and 6d, and Table S4). It should be noted that, residues in the intracellular regions (other than transducer binding) can also play a crucial role in allosteric communications, resulting in specific downstream signaling. For instance, a recent experimental and simulation study on AT1R revealed that several residues in TM4 favored *β*-arrestin signaling upon mutations, either due to enhanced *β*-arrestin recruitment or reduced G_q_protein coupling.^126^ Interestingly, in our simulations, TM4 exhibits a stronger allosteric coupling with the agonist in B2RY compared to the *β*-arrestin favoring receptor B2RTYY (Figure 6b), suggesting a similar role of TM4 residues in *β*_2_AR transducer selectivity.

Next, we compared the number of suboptimal paths from the agonist to residues at the transducer binding sites in B2RY and B2RTYY with B2RWT to understand the changes in allosteric communications that led to specific conformational rearrangements. Many hydrophobic and polar/charged residues in the intracellular part of TM5 in *β*_2_AR were found to establish contacts with the α5 helix of the G_s_protein.^51^ This region was also identified as crucial for G protein-specific signaling, as demonstrated in an experimental study that utilized pepducins (peptides derived from the intracellular end of TM5 and ICL3) as biased allosteric agents.^124^ We show the shortest paths (with thickness proportional to *n*_SOP_) from BI-167107 to the hydrophobic residues V222^5.61^ and A226^5.65^, along with Q229^5.68^ that coordinate polar interactions with α5 helix of G_s_(Figure 7; cyan). Notably, the residues V222^5.61^ and Q229^5.68^ were recognized for their high intolerance to mutation in G protein signaling response in *β*_2_AR.^127^ A significant increase in allosteric communication to all three TM5 residues in B2RY is observed with respect to B2RWT (Figures 7a and 7b; Table S3). In addition, residues in the N-terminal part of ICL3, which were experimentally observed to be involved in enhanced G protein signaling,^124^ also exhibit greater *n*_SOP_ in B2RY with respect to both B2RWT and B2RTYY (Figure S13).

**Figure 7.**
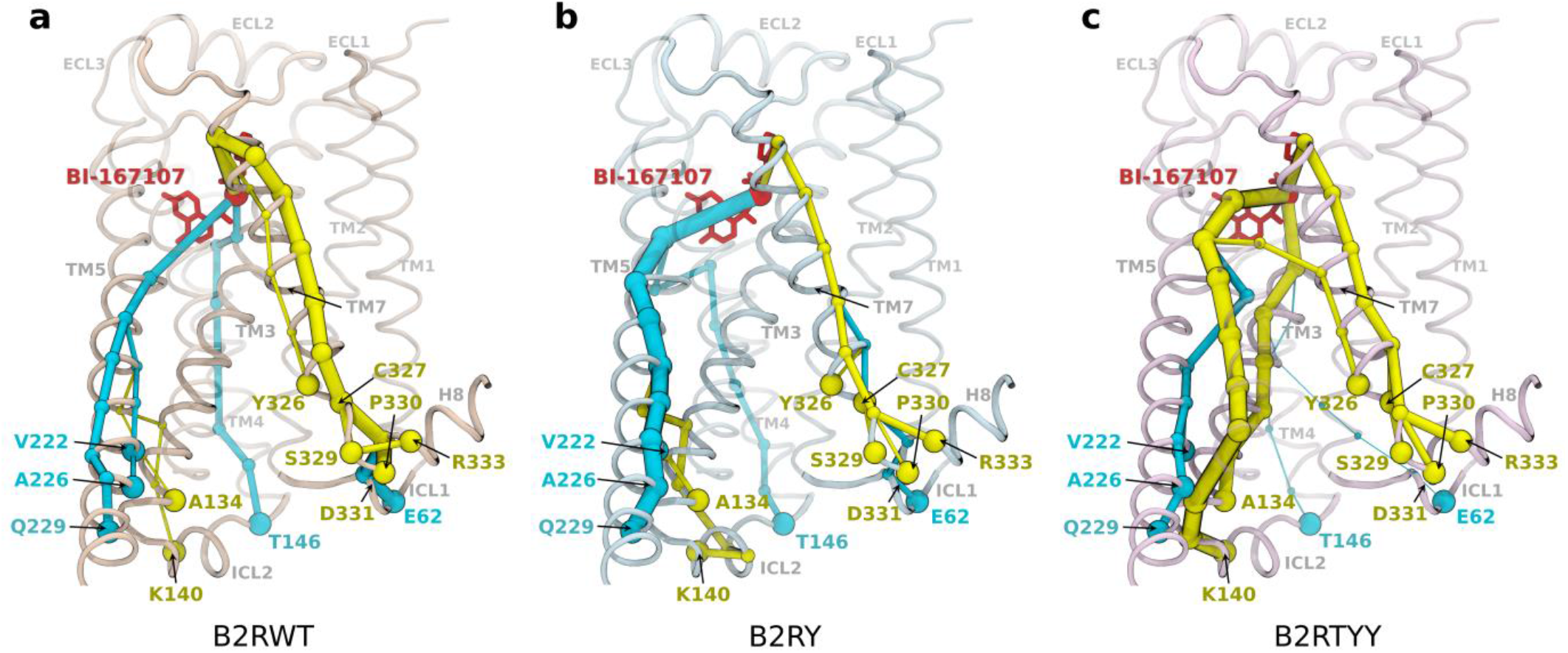
**(a-c)** Illustration of allosteric communications from the agonist BI-167107 to representative residues at the transducer interfaces of *β*_2_AR for each system; the shortest paths are used for representation, and their thickness is proportional to the number of suboptimal paths. The agonist is highlighted in red, while the paths to the G protein interface are shown in cyan, and the *β*-arrestin/GRK interface is in yellow.

Modeling the *β*-arrestin and GRK interfaces of *β*_2_AR reveals that the finger loop of *β*-arrestin and the N-terminal helix of GRK interact with H8 and TM7 (Figures 6c and 6d; Table S4). In contrast, the G protein is not directly in contact with these regions in the *β*_2_AR-G_s_complex.^51^ They were also identified as critical for the *β*-arrestin bias of *β*_2_AR using fluorine-19 nuclear magnetic resonance (^19^F NMR) spectroscopy.^43^ Furthermore, ICL2 and ICL3 that were known to interact with G protein and *β*-arrestin,^2,82,124,128^ have also been shown to involve in GRK binding using an integrated approach of cross-linking, hydrogen-deuterium exchange mass spectrometry (MS), and several other experiments.^94^ We also note that the H8 is involved in receptor-kinase interactions of rhodopsin^93^ and neurotensin receptor 1.^103^ In a cryo-EM study of the rhodopsin-GRK1 complex, the interactions of H8 residues at positions 8.48 and 8.51 with a serine residue in GRK1 (S5) were suggested to play a critical role in the phosphorylation of class A GPCRs including *β*_2_AR by GRKs.^93^ In our simulations, the residue P330^8.48^ shows a pronounced reduction in *n*_SOP_ for B2RY compared with B2RWT, consistent with its inability to bind GRK. Additional N-terminal H8 residues (S329^8.47^, D331^8.49^, and R333^8.51^) also exhibit reduced allosteric communication in B2RY, whereas both B2RWT and B2RTYY show comparable *n*_SOP_ values (Figure 7). Moreover, most ICL2 residues that constitute a common transducer interface (Figures 6c and 6d), show an increased allosteric coupling in both B2RY and B2RTYY when compared to B2RWT (Table S3).

However, this increase is prominent in B2RTYY, emphasizing their greater significance in *β*-arrestin-mediated signaling. For instance, we show the allosteric paths to K140^ICL2^ in Figure 7.

The bias against G protein signaling for B2RTYY is evident from the allosteric pathways between the agonist BI-167107 and intracellular regions. For example, the G_s_binding residue E62^ICL1^, and the TM4 residue T146^4.38^ show a substantial decrease in *n*_SOP_ in B2RTYY relative to B2RWT (Figure 7 and Table S3). Though the intracellular tip residue T146^4.38^ of *β*_2_AR is not in direct contact with G_s_, an alanine substitution for this conserved threonine yielded *β*-arrestin specificity for AT1R,^126^ suggesting the significance of T^4.38^ in biased signaling of class A GPCRs. Furthermore, the TM7 residue C327^7.54^, which exhibited notable variation in ^19^F NMR spectra upon stimulating *β*_2_AR with *β*-arrestin biased agonist,^43^ along with its neighboring residue Y326^7.53^ that interact with *β*-arrestin/GRK (Figures 6c and 6d), exhibits either comparable or enhanced allosteric signals from the agonist in B2RTYY, while a reduction is observed in B2RY, with respect to B2RWT (Figure 7 and Table S3). Moreover, allosteric signals to the residues at the intracellular end of TM3 show a similar trend as that of ICL2 (i.e., a significant increase in B2RTYY relative to the others), implying the importance of this TM3 region in *β*-arrestin selective signaling of *β*_2_AR (Table S3). For example, suboptimal pathways to the *β*-arrestin interface residue A134^3.53^ are depicted in Figure 7.

The different transducer interfaces receive allosteric communications from the agonist through various transmembrane residues. The residues that prominently feature in most suboptimal paths act as allosteric hubs and play pivotal roles in information transfer, leading to specific downstream signaling events. To identify allosteric hub residues contributing to the bias in the mutant receptors, B2RY and B2RTYY, we calculate the number of suboptimal paths passing through TM residues from the agonist to the transducer interfaces (shown in Figures 6c and 6d). The top twenty residues that most frequently appear in suboptimal paths leading to the G protein and GRK/*β*-arrestin interfaces for the wild-type are identified, and the *n*_SOP_ through these residues are compared with the mutants (Figure 8). The majority of residues (14 out of 20) display a considerable decrease in *n*_SOP_ to the GRK/*β*-arrestin interface in B2RY compared to B2RWT (Figure 8a). The reduction is primarily noticeable for the residues in TM7 (notably, Y308^7.35^, L310^7.37^, N318^7.45^, N322^7.49^, and C327^7.54^) and the mid/intracellular region of TM3 (particularly, V114^3.33^ and T118^3.37^), indicating their potential role in impairing GRK/*β*-arrestin signaling in B2RY. In fact, in an MD simulation study, residue Y308^7.35^ of TM7 was identified as a prominent contributor to the allosteric communication pathways leading to the *β*-arrestin interface than to the G protein interface.^119^ Furthermore, in our previous simulation study^44^ a significant information transfer was observed from the GRK2-phosphorylated C-terminal of *β*_2_AR to the TM3 residue T118^3.37^, suggesting its importance in GRK/*β*-arrestin signaling. Besides, in comparison to B2RWT, the number of suboptimal paths via residues to the G protein interface is substantially increased in B2RY (Figure 8b).

**Figure 8.**
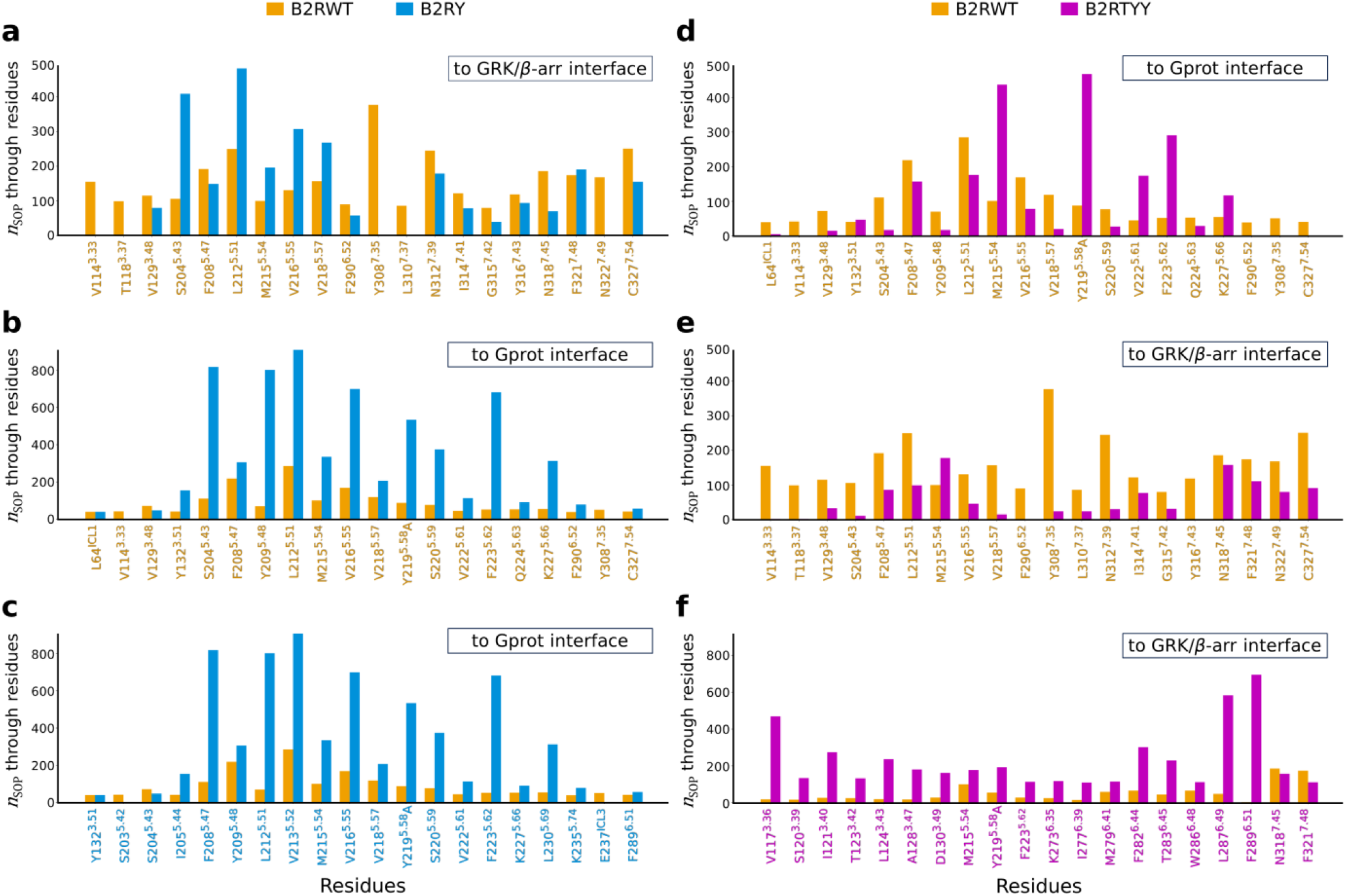
**(a-f)** Number of suboptimal paths (*n*_SOP_) passing through TM residues with BI-167107 as source and transducer (G protein and GRK/*β*-arrestin) interfaces as sinks. The top twenty residues that appear in the majority of suboptimal paths to different interfaces (marked in boxes) for the wild-type receptor B2RWT are shown in a, b, d, and e (marked in orange); *n*_SOP_ through these residues are shown as bars with colors - orange for B2RWT, blue for B2RY, and magenta for B2RTYY. The top twenty TM residues to the G protein interface for the single mutant B2RY are shown in c (marked in blue), and to the GRK/*β*-arrestin interface for the triple mutant B2RTYY in f (marked in magenta).

We also identify the residues in B2RY through which strong allosteric coupling occurs between the agonist and G protein interface (Figure 8c). Interestingly, apart from the residues that were showing increased communications (Figure 8b), a number of additional residues (8 out of 20) mediate a strong allosteric coupling with the G protein interface in B2RY (Figure 8c). The residues that act as allosteric hubs are primarily found in TM5 (S204^5.43^, Y209^5.48^, L212^5.51^, V216^5.55^, and F223^5.62^). We note that residue S204^5.43^ has also been identified as critical for G protein bias in *β*_2_AR due to its location at the base of the orthosteric pocket.^24,45,120^ This suggests Y209^5.48^, being located at the base of the pocket, could also play a role in G protein-biased signaling. In addition, as compared to the Y219^5.58^ in B2RWT, the allosteric communication via the substituted alanine is observed to be significantly enhanced to the G protein interface in B2RY (Figures 8b and 8c).

Most of the top-ranked residues (14 out of 20) involved in the allosteric signal transfer to the G protein interface in B2RWT display a notable reduction in communication strength in B2RTYY (Figure 8d). The reduction is primarily observed for the residues in TM5 (notably, F208^5.47^, Y209^5.48^, L212^5.51^, V216^5.55^, and V218^5.57^). Strikingly, the signal transfer through the TM6 residue F290^6.52^, and the TM7 residues Y308^7.35^ and C327^7.54^ to the G protein interface is entirely disrupted in B2RTYY (Figure 8d). However, as opposed to intuition, allosteric communications through the residues that transmit information to the GRK/*β*-arrestin interface in B2RWT are diminished in B2RTYY (Figure 8e). Nonetheless, due to a significant rearrangement of the allosteric network in B2RTYY, a different set of residues appears to act as strong allosteric hubs between the agonist and the GRK/*β*-arrestin interface (Figure 8f), where primarily they reside in the mid/intracellular region of TM3 (V117^3.36^, I121^3.40^, L124^3.43^, and D130^3.49^) and the mid/extracellular region of TM6 (F282^6.44^, T283^6.44^, W286^6.48^, L287^6.49^, and F289^6.51^). In B2RTYY, the appearance of D130^3.49^ as one of the allosteric hub residues and an increased preference for the downward rotameric state of residue R131^3.50^ suggest the involvement of the conserved D(E)RY motif in *β*-arrestin signaling bias of *β*_2_AR. We further note that consistent with our findings, the TM6 residues F282^6.44^, W286^6.48^, and F289^6.51^ also appeared in the allosteric paths from the *β*-arrestin-biased agonist ethylnorepinephrine to the intracellular regions in a simulation study of *β*_2_AR.^33^

It is well known that the perturbations, including ligand/transducer binding,^26,33,34,37,40,86,121^ post-translational modifications such as phosphorylation,^44,129^ and mutation,^19^ trigger conformational changes leading to activation and biased signaling of GPCRs. The distinct reconfigurations of allosteric pathways, connecting the extracellular agonist and different transducer interfaces, due to mutations, are suggestive of driving specific conformations of *β*_2_AR that supports the observed bias in experiments.^17,18^

## Conclusion

The phenomenon of receptor bias refers to the selective signaling through a GPCR, mediated by either G protein or *β*-arrestin, due to mutations. Investigating the functional dynamics of biased receptors holds significant potential in elucidating the fundamentals of transducer selective signaling in GPCRs induced by biased ligands. In *β*_2_AR, a single mutation (Y219^5.58^A) selects against GRK binding, thus, disfavoring the association with *β*-arrestins, leading to G protein bias of the receptor.^18^ Whereas, a triple mutation (T68^2.39^F, Y132^3.51^G, and Y219^5.58^A) inhibits G protein signaling, consequently resulting in *β*-arrestin bias.^17^ In an attempt to understand the atomistic basis of this biased signaling in *β*_2_AR, we performed large-scale all-atom enhanced sampling GaMD simulations and analyzed the allosteric conformational changes sampled by mutant receptors.

Our observations reveal specific conformations of the transmembrane helices in the extracellular and cytosolic regions of the mutant receptors compared to the wild-type. In the single mutant, ICL3 mostly positions away from the transducer-binding cavity, sampling an open state that facilitates G protein binding. In contrast, the triple mutant prefers a closed state of the ICL3, thereby occluding the cavity for G protein engagement. Further, in the mutant receptors, the side chains of R131^3.50^ and Y326^7.53^ in the conserved motifs D(E)RY and NPxxY exhibit characteristic orientations that could enable specific transducer interactions. In particular, the *β*-arrestin-favoring triple mutant displays a relatively larger population of the downward rotameric state of R131^3.50^, characterized by a cytoplasm-facing side chain conformation that selectively facilitates the engagement of *β*-arrestin over G protein. By employing machine learning classification algorithms, we discern the inter-residue interactions that promote different orientations of R131^3.50^ and Y326^7.53^ in the wild-type and mutant systems. The evaluation of suboptimal paths reveals distinctive rewiring of allosteric communication pathways between the extracellular agonist BI-167107 and the residues in the interfaces of various transducers (G protein and GRK/*β*-arrestin). These allosteric reconfigurations drive specific conformational sampling, resulting in the selective engagement of transducers in both single and triple mutants. The critical residues identified in allosteric signal transfer for each *β*_2_AR mutant present promising opportunities for future experimental and computational investigations that could target the sites to unravel the allosteric mechanisms underlying biased signaling across various other GPCRs. Moreover, the atomistic insights presented here may help in developing therapeutic strategies for diseases caused by mutations in GPCRs and aid in designing biased drugs that target these receptors with fewer side effects and better efficacy.

## Associated Content

### Data Availability Statement

To identify inter-residue contacts, we have created a Python program called *trajcontacts*, which can be accessed at https://github.com/rkmlabiiserb/trajcontacts. All the essential input files required to reproduce the simulations and machine learning analysis codes are available at https://github.com/rkmlabiiserb/b2armut.

## Supporting Information

Additional methods details: model preparation and system setup, MD simulation protocols, theory of Gaussian-accelerated MD simulation, ML-based methods, contact analysis, and theory of generalized correlations based on mutual information are given as supplementary methods.

Figure S1: schematic representation of the *β*_2_AR systems considered for simulations. Figure S2: BIC scores obtained in the RMSD-based clustering using GMM. Figure S3: representative structures from clusters of all systems not shown in Figure 1. Figures S4-S6: distributions of the intracellular displacement of TM6, the intracellular shift of TM5 from the receptor core, and backbone RMSD of the NPxxY motif. Figure S7: representation of the average volume of transducer-binding cavities and distribution plots of intracellular TM3-TM6 distances and H8 displacement. Figure S8: cartoon representations showing D(E)RY and NPxxY motif in the active and inactive *β*_2_AR and orientation of side chains of R131^3.50^ and Y326^7.53^ in various experimental structures. Figure S9: silhouette coefficient scores obtained in the 2D *k*-means clustering of χ1 torsions of R131^3.50^ and Y326^7.53^. Figure S10: nonbonding interaction energies of R131^3.50^ and Y326^7.53^ with mutated residues and time series representation of water-mediated interaction between Y219^5.58^ and Y326^3.26^ in B2RWT as well as N-O distance between R131^3.50^ and E268^6.30^. Figure S11: difference in agonist interaction energies between mutant and wild-type systems. Figure S12: heatmap of suboptimal paths with *l*_cut_= 15 for each system. Figure S13: representation of allosteric communication paths to ICL3.

Table S1: summary of GaMD simulations conducted for each *β*_2_AR system. Table S2: performance evaluation metrics of ML-based classifications. Table S3: number of suboptimal paths from the agonist to the residues at the intracellular regions for all systems calculated using *l*_cut_ = 10 and 15. Table S4: list of *β*_2_AR residues that are in the interface of G protein/GRK/*β*-arrestin.

References 130 to 139 are in the Supporting Information. ^130–139^

## Author Information

## Author Contributions

R.K.M. designed and supervised the research. M.K.M. carried out MD simulations, analyzed the data, and prepared the manuscript. M.K.M., K.S., and R.K.M. edited and approved the final version of the manuscript.

## Notes

The authors declare no competing financial interest.

## Acknowledgment

M.K.M. was supported by a research fellowship granted by the Council of Scientific & Industrial Research (CSIR), India. K.S. was supported by the research fellowships provided by IISER Bhopal. R.K.M. acknowledges the financial support from the Science and Engineering Research Board (SERB), Department of Science and Technology, India (File No. EMR/2016/006815).

---

“For Table of Contents Only”

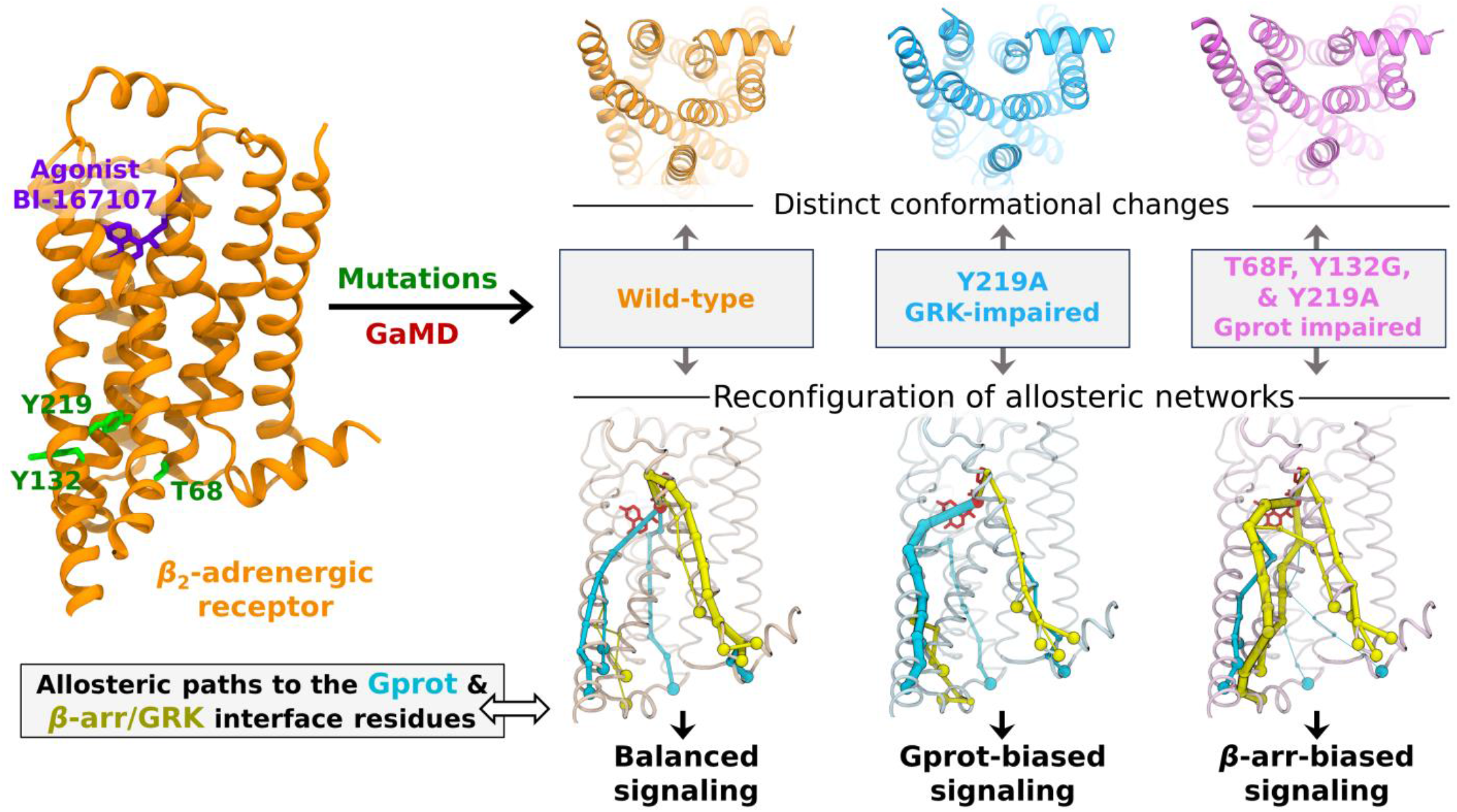

## Notes

### Competing Interest Statement

The authors have declared no competing interest.

