## Supplemental File for "Biased Signaling in Mutated Variants of *β*_2_-Adrenergic Receptor: Insights from Molecular Dynamics Simulations"

AUTHOR INFORMATION

*__Corresponding Author__

### Supplementary Methods

#### Model preparation and system setup

All the missing parts except the first 28 residues at the N-terminus and the last 72 residues at the C-terminus of the $\beta_2$AR were modeled after reverting the engineered mutations (T96$^{2.67}$M, T98$^{ECL1}$M, and E187$^{ECL3}$N) in the wild-type structure (PDB ID: 3SN6) using MODELLER v9.17.[1] Missing residues in the extracellular loop 2 (ECL2) were modeled using the inactive crystal structure (PDB ID: 2RH1) as the template.[2] The intracellular loop 3 (ICL3) residues (F240$^{ICL3}$ to S261$^{ICL3}$) and three residues at the tip of TM6 (S262$^{6.24}$ to F264$^{6.26}$) were modeled using the secondary structure information predicted using Jpred 4 and PSIPRED.[3,4] The dominant protonation states of all titrable solvent-exposed residues were maintained, and residues D79$^{2.50}$, E122$^{3.41}$, D130$^{3.49}$, and H172$^{4.64}$ were protonated.[5] The disulfide bonds resolved in the crystal structure (C106$^{3.25}$ - C191$^{ECL2}$ and C184$^{ECL2}$ - C190$^{ECL2}$) were retained for the simulations. To maintain the salt concentration of 150 mM and to neutralize the system, 67 sodium and 46 chloride ions were added. The final wild-type system in a rectangular simulation box contained 67,579 atoms, including 148 lipids and 14,173 water molecules, and initially measured roughly 79×79×116 Å$^3$ in volume. The single and triple mutant systems had similar box dimensions and approximately the same number of atoms as in the wild-type.

#### Molecular dynamics simulations details

Prior to the conventional MD (cMD) equilibrations, the system underwent an initial energy minimization process in three steps. Each step involved 2500 cycles of steepest descent followed by 2500 cycles of the conjugate gradient method. During these energy minimization steps, we applied positional restraints with harmonic force constants as follows: (i) 10.0 kcal mol$^{-1}$ Å$^{-2}$ on all heavy atoms of the protein and phosphorous atoms of lipids and dihedral restraints of 250 kcal mol$^{-1}$ rad$^{-2}$ on dihedral angles of lipids, (ii) 5.0 kcal mol$^{-1}$ Å$^{-2}$ on all heavy atoms of the protein and phosphorous atoms of lipids and dihedral restraints of 100 kcal mol$^{-1}$ rad$^{-2}$ on dihedral angles of lipids, and (iii) 1.0 kcal mol$^{-1}$ Å$^{-2}$ on all heavy atoms of the protein and phosphorous atoms of lipids. Next, we gradually heated the system to 310 K in two steps, employing a 1 fs time step and maintaining positional restraints and dihedral restraints: (i) 0 to 100 K in the NVT ensemble for 12.5 ps, and (ii) 100 to 310 K in the NPT ensemble of 125 ps. The system was further equilibrated in the NPT ensemble with a 2 fs time step, gradually reducing the dihedral restraints on lipids stepwise (100, 50, and 25 kcal mol$^{-1}$ rad$^{-2}$), and subsequently reducing the positional restraints stepwise (7.5, 5.0, 2.5, 1.0, 0.5, and 0.1 kcal mol$^{-1}$ Å$^{-2}$), with each step involving a 1 ns simulation until no restraints remained on the lipid bilayer. The lipids were further equilibrated in the presence of solvent (water and ions), keeping the protein restrained for 100 ns. The restraint on heavy atoms of protein was then removed gradually by reducing the harmonic force constants (10, 7.5, 5.0, 4.0, 3.0, 2.0, 1.0, 0.9, 0.8, 0.7, 0.6, 0.5, 0.4, 0.3, 0.2, and 0.1 kcal mol$^{-1}$ Å$^{-2}$), with each step simulated for 5 ns. The Langevin thermostat was employed to maintain the temperature, while the Monte Carlo barostat was used to keep the pressure at 1 bar throughout the simulations. The SHAKE algorithm was applied to constrain bonds involving hydrogen atoms, and the SETTLE algorithm was used for water molecules. Periodic boundary conditions were utilized, and a cut-off of 12 Å was set for nonbonded interactions, with a force-switch applied from 10 Å. The particle mesh Ewald (PME) method was used to calculate long-range electrostatic interactions, employing a cubic B-spline interpolation of order 4 and implicitly computing the Ewald coefficient. Further, the system was equilibrated for 100 ns, and the last frame was extracted for mutations.

While preparing the triple mutant system, 25 ns of cMD simulations were performed after mutating each residue to attain local equilibration around mutated residues. Lastly, an additional 100 ns cMD equilibration simulations were performed for both the mutated systems in the NPT ensemble.

### Gaussian-accelerated MD simulation

In GaMD, a harmonic boost potential, $\Delta V(\boldsymbol{r})$, is added to the system potential, $V(\boldsymbol{r})$, to overcome the energy barrier,[6,7]

$$V^*(\boldsymbol{r}) = V(\boldsymbol{r}) + \Delta V(\boldsymbol{r}),$$

$$\Delta V(\boldsymbol{r}) = \begin{cases} \frac{1}{2}\kappa(\varepsilon - V(\boldsymbol{r}))^2, V(\boldsymbol{r}) < \varepsilon \\ 0, V(\boldsymbol{r}) \geq \varepsilon \end{cases}, \tag{1}$$

where $V^*(\boldsymbol{r})$ is the modified system potential, $\boldsymbol{r} = \{\boldsymbol{r_1}, \boldsymbol{r_2}, \dots, \boldsymbol{r_N}\}$ is the coordinate vector of $N$ atoms in the system, $\kappa$ is the harmonic force constant, and $\varepsilon$ is the threshold cut-off for $V(\boldsymbol{r})$, below which the boost potential is added. The value of $\varepsilon$ is set in the range,

$$V_{max} \leq \varepsilon \leq V_{min} + \frac{1}{\kappa}, \tag{2}$$

where $V_{max}$ is the maximum and $V_{min}$ is the minimum of system potential energies, and the value of $\kappa$ must be $\kappa \leq \frac{1}{V_{max}-V_{min}}$. If $\kappa$ is defined as, $\kappa_0 \frac{1}{V_{max}-V_{min}}$, then $\kappa_0$ is in the range of $0 < \kappa_0 \leq 1$. For accurate energetic reweighting, the distribution of boost potential $\Delta V$ should be narrow enough (having a small standard deviation) such that, $\sigma_{\Delta V} = \kappa(\varepsilon - V_{avg})\sigma_V \leq \sigma_0$, where $V_{avg}$ and $\sigma_V$ denote the average and standard deviation of $V$, and $\sigma_0$ is the user-defined upper limit ($\text{e.g.}, 10k_BT$) of $\sigma_{\Delta V}$.[8] When the threshold energy $\varepsilon = V_{max}$ (lower bound of $\varepsilon$), $\kappa_0$ can be calculated as,[6,7]

$$\kappa_0 = \min(1, \kappa_0') = \min\left(1, \frac{\sigma_0}{\sigma_V} \cdot \frac{V_{max} - V_{min}}{V_{max} - V_{avg}}\right) \tag{3}$$

Alternatively, for $\varepsilon = V_{min} + 1/\kappa$ (upper bound of $\varepsilon$),

$$\kappa_0 = \kappa_0'' \equiv \left(1 - \frac{\sigma_0}{\sigma_V}\right)\frac{V_{max} - V_{min}}{V_{avg} - V_{min}}, \tag{4}$$

where $\kappa_0''$ is obtained in the range of 0 to 1; otherwise, $\kappa_0$ is estimated using equation 3.

For the free energy calculations, the canonical ensemble probability distribution ($p(A)$) along any reaction coordinate $A(\boldsymbol{r})$ was recovered by reweighting its biased (GaMD) probability distribution $p^*(A)$, where $\boldsymbol{r}$ denotes the atomic positions, $\boldsymbol{r} = \{\boldsymbol{r_1}, \boldsymbol{r_2}, \dots, \boldsymbol{r_N}\}$ and $N$ denotes the total number of atoms. Using the boost potential $\Delta V(\boldsymbol{r})$ for each simulation frame, $p(A)$ is calculated as,[6,7]

$$p(A_i) = p^*(A_i)\frac{\left\langle e^{\beta \Delta V(r)}\right\rangle_i}{\sum_{j=1}^{M}\left\langle p^*(A_j)e^{\beta \Delta V(r)}\right\rangle_j}, i = 1, 2, \dots, M, \tag{5}$$

where $M$ is the number of bins and $\left\langle e^{\beta \Delta V(r)}\right\rangle_i$ is the ensemble-averaged reweighting factor of $\Delta V(\boldsymbol{r})$ for simulation frames in the $i^{\text{th}}$ bin, where $\beta = 1/k_BT$ and $k_B$ is the Boltzmann constant. The reweighting factor is approximated using the cumulant expansion,

$$\langle e^{\beta \Delta V} \rangle = exp\left\{\sum_{k=1}^{\infty} \frac{\beta^k}{k!} C_k\right\} \quad (6)$$

Usually, $\Delta V(\boldsymbol{r})$ follows near-Gaussian distribution, and thus, cumulant expansion to the second order can be a good approximation for reweighting the distribution, where the first two cumulants are given by $C_1 = \langle \Delta V \rangle$ and $C_2 = \langle \Delta V^2 \rangle - \langle \Delta V \rangle^2 = \sigma_{\Delta V}^2$. The reweighted free energy or the potential of mean force (PMF) is calculated as $F(A_i) = -(1/\beta) \ln p(A_i)$.

**RMSD-based conformational clustering using Gaussian mixture model**

The Gaussian mixture model (GMM) is a probabilistic model that assumes the observed data are comprised of subpopulations, each characterized by a Gaussian distribution.[9] The algorithm estimates the parameters of these Gaussian distributions, such as their means, covariances, and weights, to model the underlying subpopulations. GMM implemented in the SciPy Python library was used for one-dimensional clustering of the root mean square deviations (RMSD) of $C_\alpha$ atoms in TM helices and H8, and the number of optimal clusters for each system was determined using the Bayesian information criterion (BIC).[10]

**2-D clustering of dihedral angels using *k*-means clustering**

The *k*-means clustering algorithm partitions a given set of data points into *k* clusters, where *k* is a predefined number.[9] It works by iteratively assigning each data point to the closest centroid, which is the mean of the data points in the cluster. After all data points are assigned to the closest centroid, the algorithm updates the centroids by re-estimating the mean of each cluster. This process continues until the centroids no longer change or a maximum number of iterations is reached. We used *k*-means implemented in the SciPy Python library for two-dimensional clustering of $\chi 1$ torsional angles of R131$^{3.50}$ and Y326$^{7.53}$ and optimized the number *k* using the silhouette coefficient method.[10]

**Feature importance analyses using classification models**

The electrostatic and van der Waals interaction energies between residue pairs involving R131$^{3.50}$ or Y326$^{7.53}$, with an average energy difference in energy $|\langle E \rangle| \geq 1$ kcal/mol, were extracted using the *CPPTRAJ* module of AMBER18 and considered as features for classification for each receptor system (3 classes for the wild-type and 4 each for the single and triple mutants, as shown in Table 1, main text). To estimate the feature importance, the data were fitted against class labels using gradient boosting, extreme gradient boosting (XGBoost), random forest, and the automated approach AutoGluon developed by Amazon Web Services.[11]

For assessing the model performance, we utilized log loss (cross-entropy loss) in addition to the commonly used accuracy metric, as the former is effective in evaluating imbalanced class distributions. Accuracy measures the proportion of correctly classified instances out of all predictions, calculated as,

$$\text{accuracy} = \text{Number of correct predictions/Total number of predictions}$$

Log loss is a metric that penalizes false classifications based on predicted probabilities for each class, estimated as,

$$\text{log loss} = -\frac{1}{N}\sum_{i=1}^{N}\sum_{i=1}^{M} y_{ij} \log(p_{ij}),$$

where $N$ is the number of instances, $M$ is the number of classes, $y_{ij}$ is the indicator function (1 if the instance $i$ belongs to class $j$, 0 otherwise), and $p_{ij}$ is the predicted probability of instance $i$ belonging to class $j$. Table S2 lists the accuracy and log loss scores obtained for each trained model.

**Identification of $\beta_2$AR residues at the interface of transducers**

The $\beta_2$AR residues at the interfaces of transducers (G protein, GRK, and $\beta$-arrestin) were identified utilizing various PDB structures using a Python script – *trajcontacts* (https://github.com/rkmlabiiserb/trajcontacts). G protein interface residues were taken from the active state $\beta_2$AR structure bound to $G_s$ (PDB ID: 3SN6).[12] Due to the unavailability of $\beta_2$AR structures complexed GRK or $\beta$-arrestin, all available GRK and $\beta$-arrestin- bound structures of class A GPCRs (PDB ID of rhodopsin-GRK1 complex is 7MTA[13] and PDB IDs of GPCR-$\beta$-arrestin 1 structures are $\beta_1$AR: 6TKO,[14] $M_2$R: 6U1N,[15] NTSR1: 6UP7 and 6PWC,[16,17] and 5-$HT_{2B}$R: 7SRS[18]) were considered for determining interface residues. The GRK interface residues were taken from aligning the $\beta_2$AR structure with 7MTA and a recent study that determined the cryo-EM structure of the NTSR1-GRK2 complex.[19] $\beta$-arrestin interface residues were determined by aligning the $\beta_2$AR structure with 6TKO, 6U1N, 6UP7, 6PWC, and 7SRS. The $\beta_2$AR residues within 4 Å proximity of transducers are listed as interface residues (Table S4).

**Generalized correlations based on mutual information**

The correlations between residue mosnomer fluctuations are frequently used to construct the allosteric network of macromolecular systems.[20,21] However, the commonly used Pearson correlation coefficient fails to capture nonlinear correlations; generalized correlations derived from mutual information (MI) provide a better measure to quantify allosteric communications between monomers.[21–23]

MI between fluctuations of $C_\alpha$ atoms in a pair of protein monomers (amino acid residues) X and Y is calculated as,[22]

$$I_{\mathrm{XY}} = H(\mathbf{X}) + H(\mathbf{Y}) - H(\mathbf{X},\mathbf{Y}), \tag{7}$$

where the vectors **X** and **Y** represent cartesian coordinates of the $C_\alpha$ atoms of the residues X and Y, $H(\mathbf{X}) = -\int p(\mathbf{X}) \ln p(\mathbf{X}) d\mathbf{X}$ and $H(\mathbf{X},\mathbf{Y}) = -\iint p(\mathbf{X},\mathbf{Y}) \ln p(\mathbf{X},\mathbf{Y}) d\mathbf{X} d\mathbf{Y}$ are the Shannon entropies, and $p(\mathbf{X})$ is the marginal probability distribution of **X** and $p(\mathbf{X},\mathbf{Y})$ is the joint probability distribution of **X** and **Y**.

An efficient estimation of Shannon entropy is given by the nearest neighbor method that exploits the statistics of the distances between neighboring data points.[24,25] The $k$-nearest neighbor ($k$-NN) technique introduced by Kraskov-Stögbauer-Grassberger (KSG) estimates MI between X and Y using the equation,[25,26]

$$I_{\mathrm{XY}} = \psi(k) - 1/k - \langle \psi(n_{\mathrm{X}}) - \psi(n_{\mathrm{Y}}) \rangle + \psi(\mathrm{N}) \tag{8}$$

where, $\psi$ is the digamma function, the angle brackets denote averaging over different time points, N is the total number of simulation frames, and $n_{\mathrm{X}}$ is the number of frames in which the position of X is close to its position in the reference frame. Parameter $k$ was set to be 6, as proposed by Lange and Grubmüller.[22,26]

### Supplementary Figures

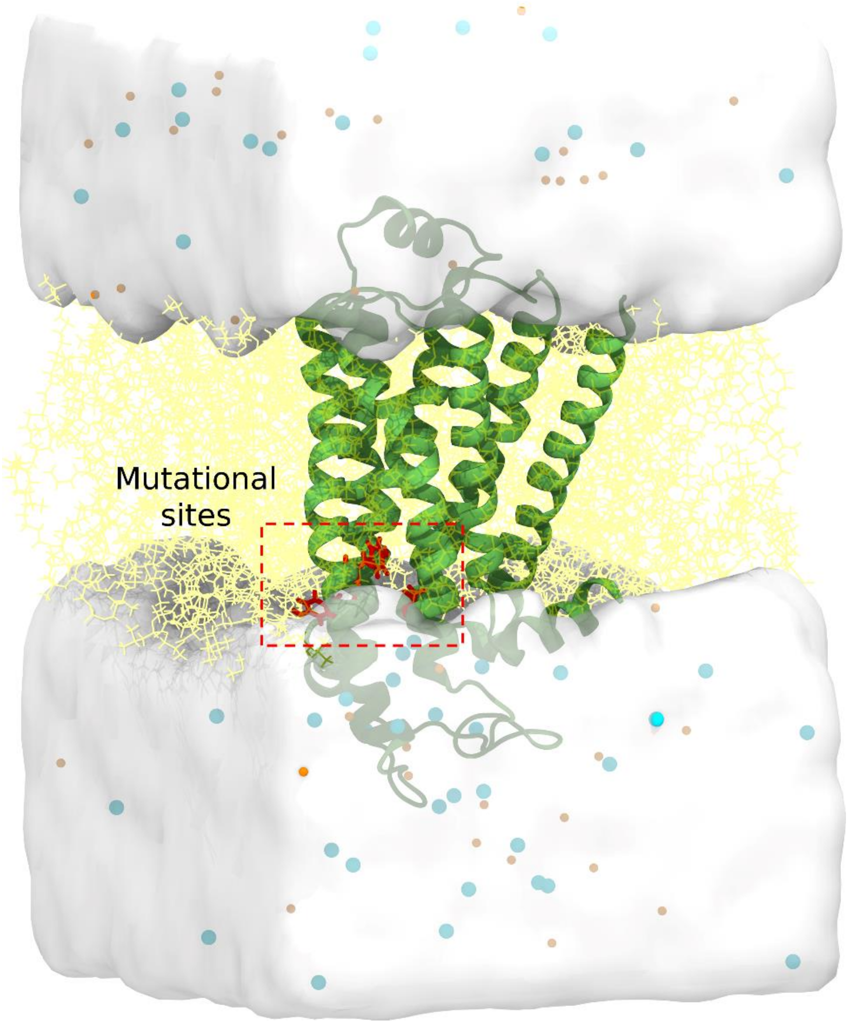


**Figure S1.** Schematic representation of $\beta_2$AR (lime green ribbon) inserted into POPC bilayer (yellow lines) and solvated in water (white transparent surface) neutralized by 150 mM sodium (orange spheres) and chloride (cyan spheres) ions.

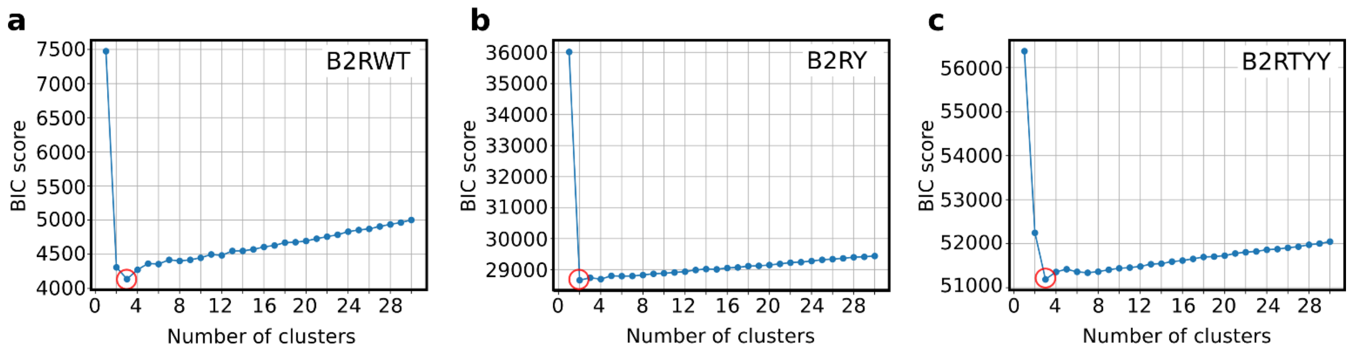


**Figure S2. (a-c)** Bayesian information criterion (BIC) scores obtained in the one-dimensional clustering of RMSD of $C_\alpha$ atoms of TM helices and H8 using the Gaussian mixture model. Minima represents the optimum number of clusters (circled in each plot).

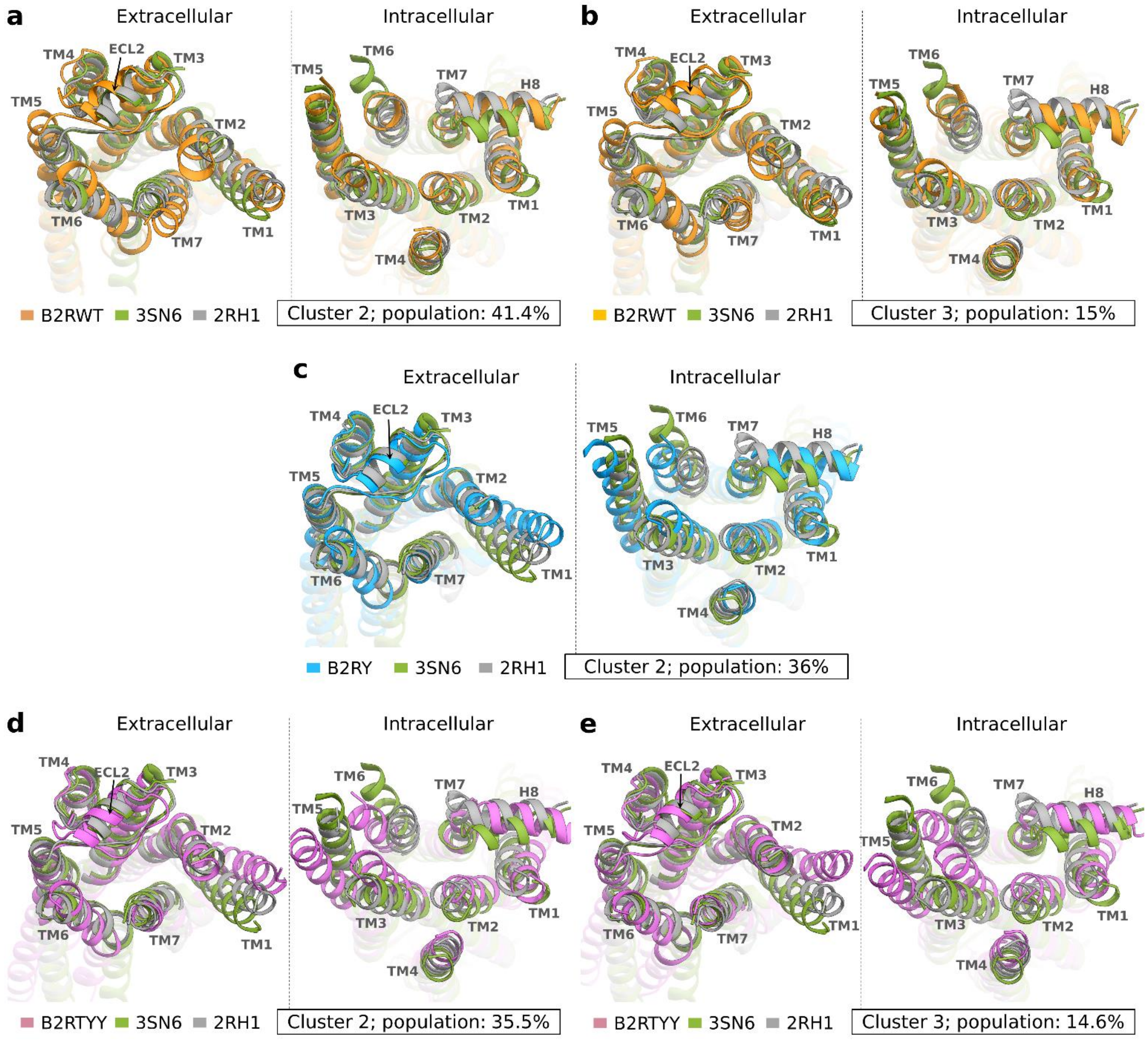


**Figure S3.** Conformational variations in the extracellular and intracellular regions of $\beta_2$AR due to point mutations identified using RMSD-based clustering. Representative snapshots extracted from cluster 2 of B2RWT, B2RY, and B2RTYY, as well as cluster 3 of B2RWT and B2RTYY, are superimposed with the active and inactive structures (PDB ID: 3SN6 and 2RH1). Cluster numbers, along with their populations, are given inside boxes.

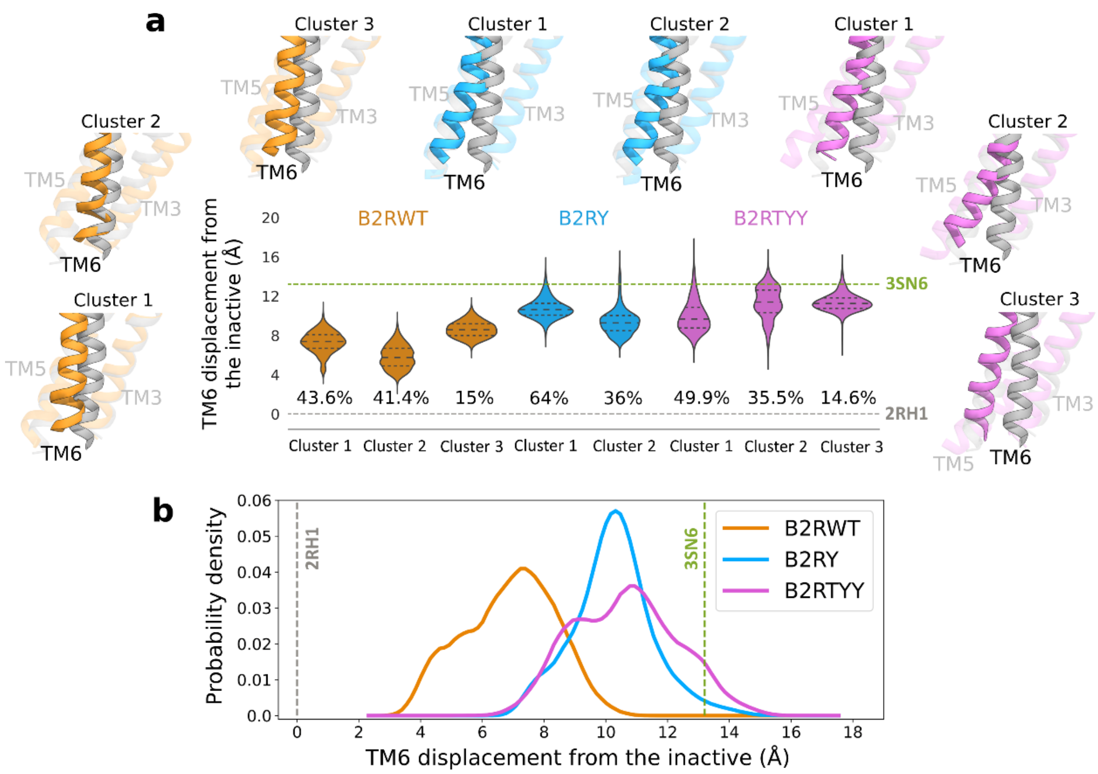


**Figure S4.** Distribution of the intracellular displacement of TM6 with respect to the inactive structure (PDB ID: 2RH1) measured at E268$^{6.30}$ $C_{\alpha}$. **(a)** Cluster-wise violin plots and **(b)** probability density plots over the entire dataset for each system.

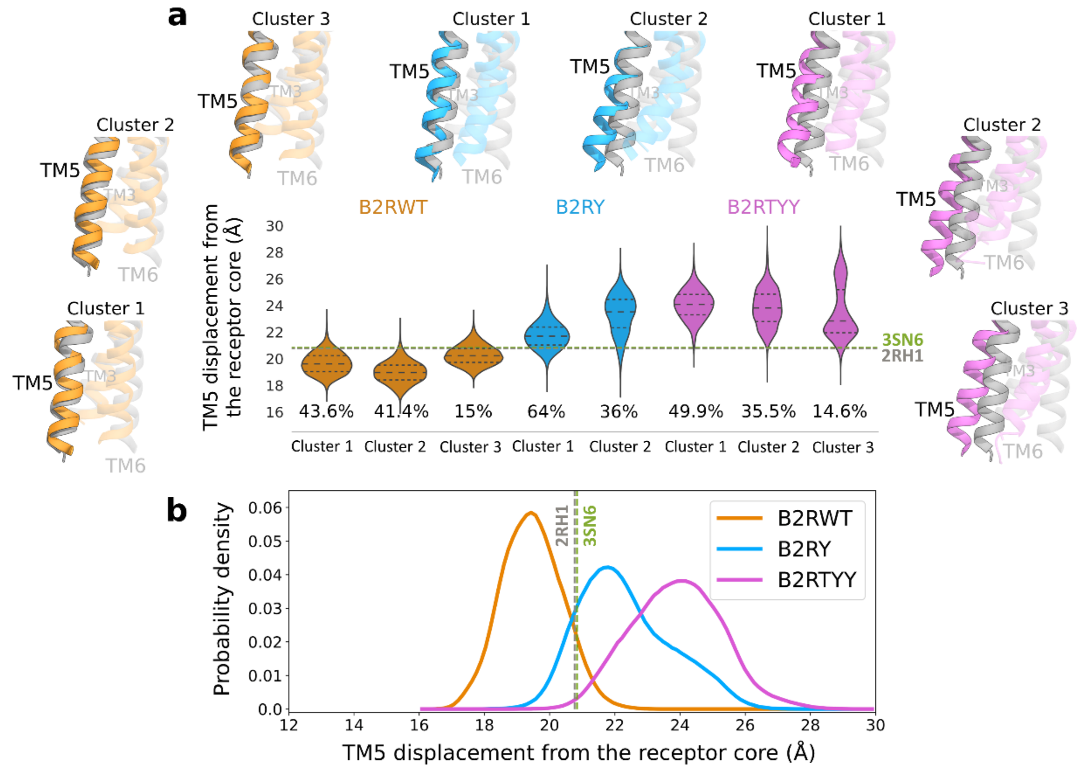


**Figure S5.** Distribution of the intracellular shift of TM5 from the receptor core (defined as the distance between L230$^{5.69}$ $C_{\alpha}$ and the geometric center of the group of $C_{\alpha}$ atoms of residues at the intracellular tip of transmembrane helices in the active state structure, PDB ID: 3SN6). **(a)** Cluster-wise violin plots and **(b)** probability density plots over the entire dataset for each system.

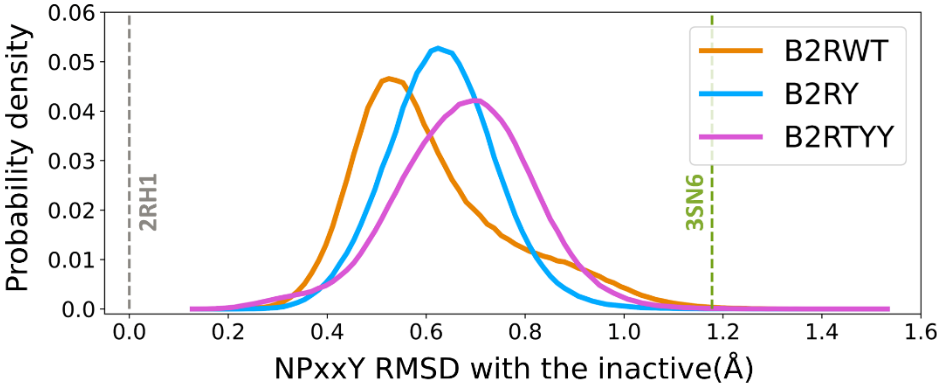


**Figure S6.** Distribution of the backbone RMSD of the NPxxY motif with respect to the inactive structure (PDB ID: 2RH1).

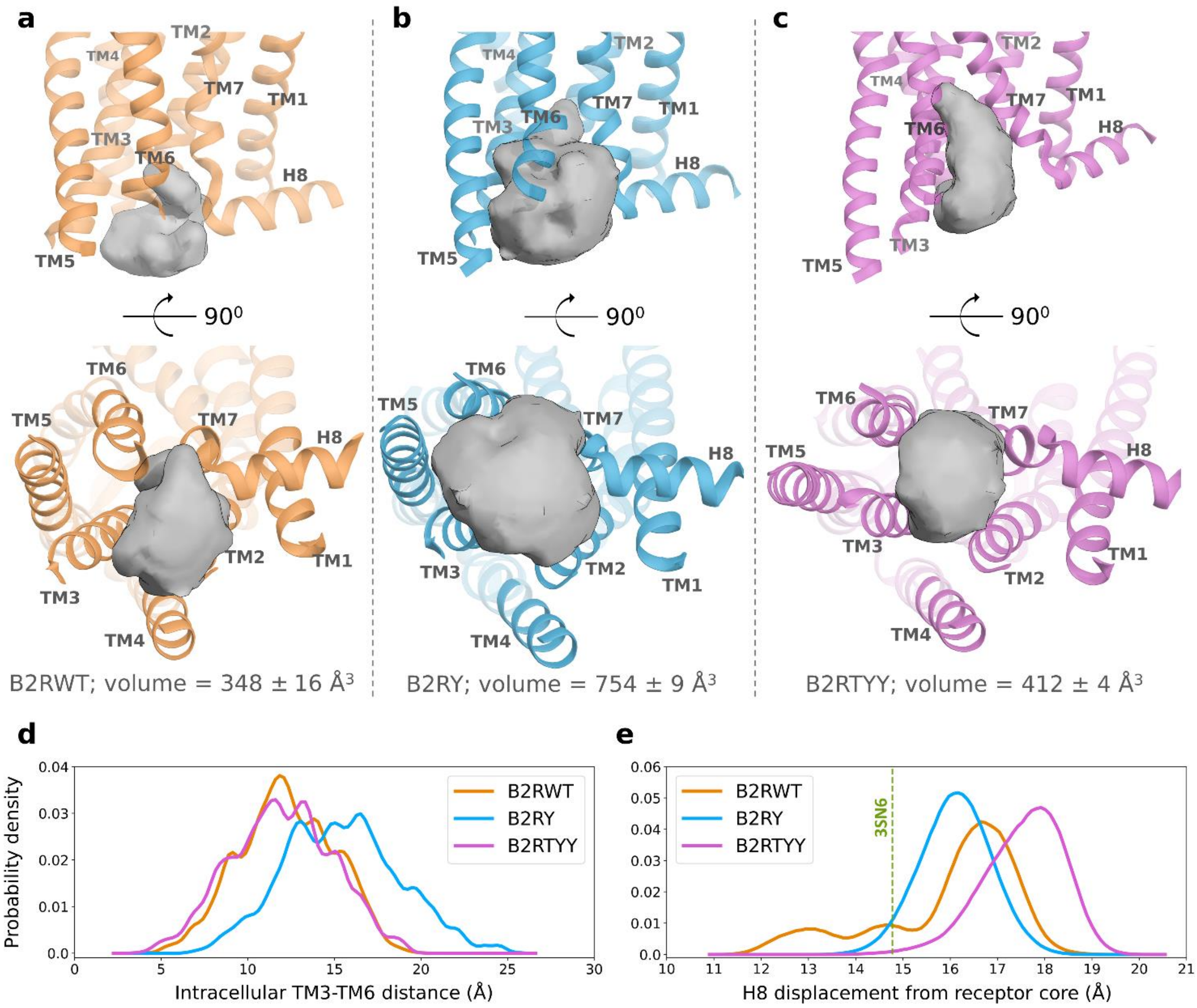


**Figure S7. (a-c)** The average volume of intracellular transducer binding cavity of each system; errors represent the standard error of mean over three trajectories. **(d)** Distribution of $C_\alpha$ distances between the intracellular tip residues of TM3 and TM6. **(e)** Distribution of H8 displacement measured between residue R333$^{8.51}$ of H8 and receptor core (defined as the geometric center of the group of $C_\alpha$ atoms of residues at the intracellular tip of transmembrane helices in the active state structure, PDB ID: 3SN6).

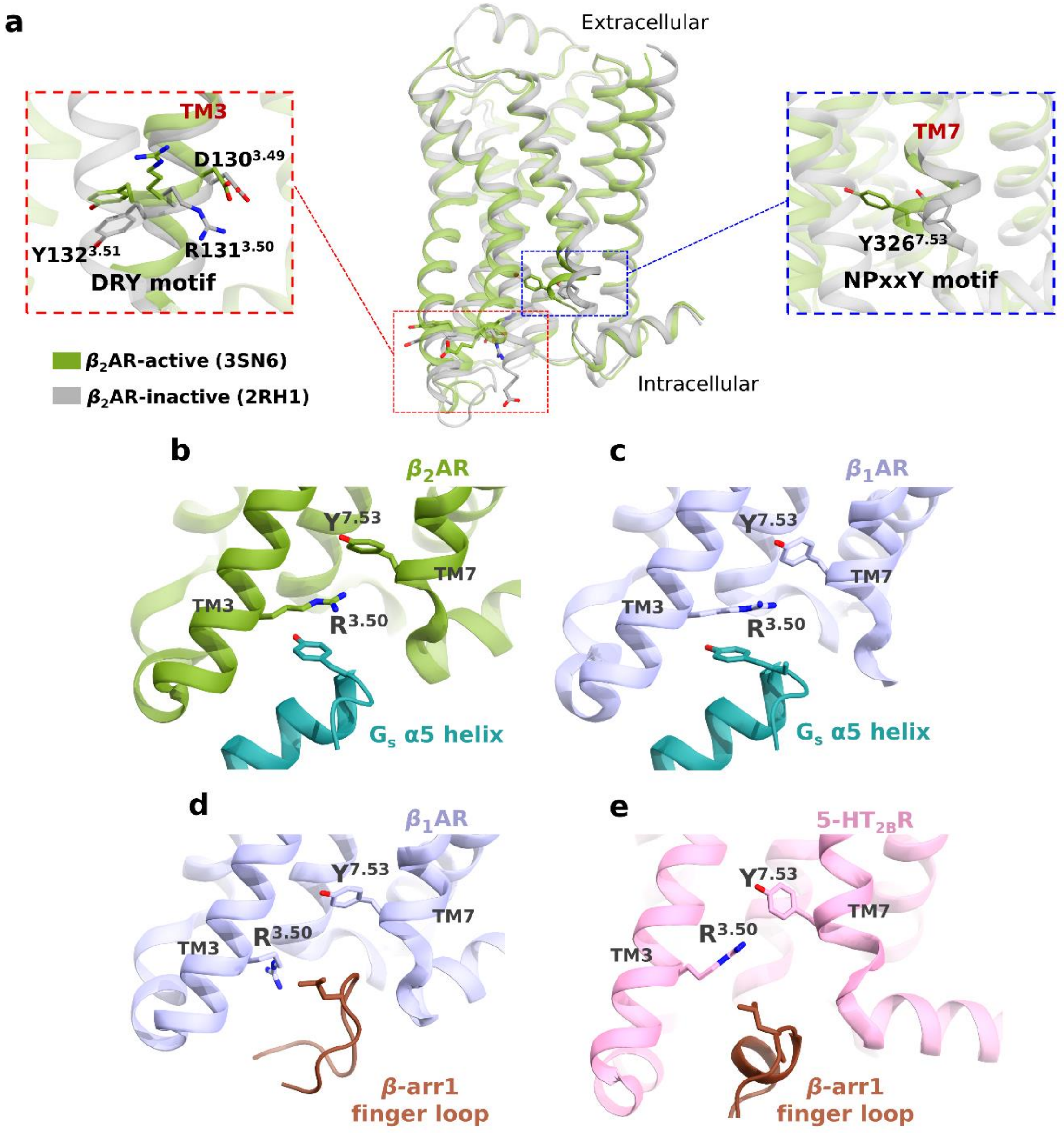


**Figure S8. (a)** The active and inactive state structures of $\beta_2$AR (PDB IDs: 3SN6 and 2RH1) marked with key conserved motifs at the intracellular region - D(E)RY, and NPxxY. **(b-e)** Orientations of $R^{3.50}$ and $Y^{7.53}$ in $\beta_2$AR-$G_s$ (PDB ID: 3SN6), $\beta_1$AR-$G_s$ (PDB ID: 7JJO), $\beta_1$AR-$\beta$arr1 (PDB ID: 6TKO), and 5-$HT_{2B}$R-$\beta$arr1(PDB ID: 7SRS) complexes.

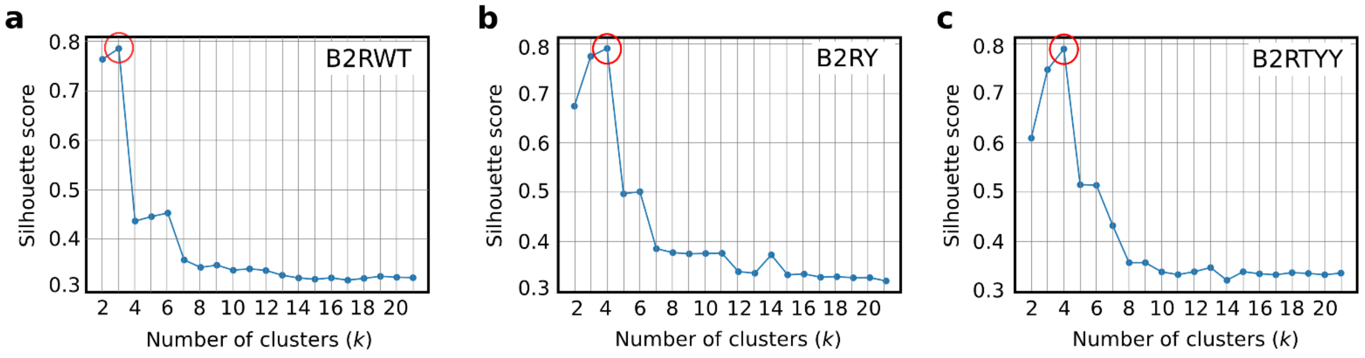


**Figure S9. (a-c)** Silhouette coefficient scores obtained in the two-dimensional *k*-means clustering of $\chi1$ torsions of $R131^{3.50}$ and $Y326^{7.53}$. Maxima represents the optimum number of clusters or optimal value of the parameter *k* (circled in each plot).

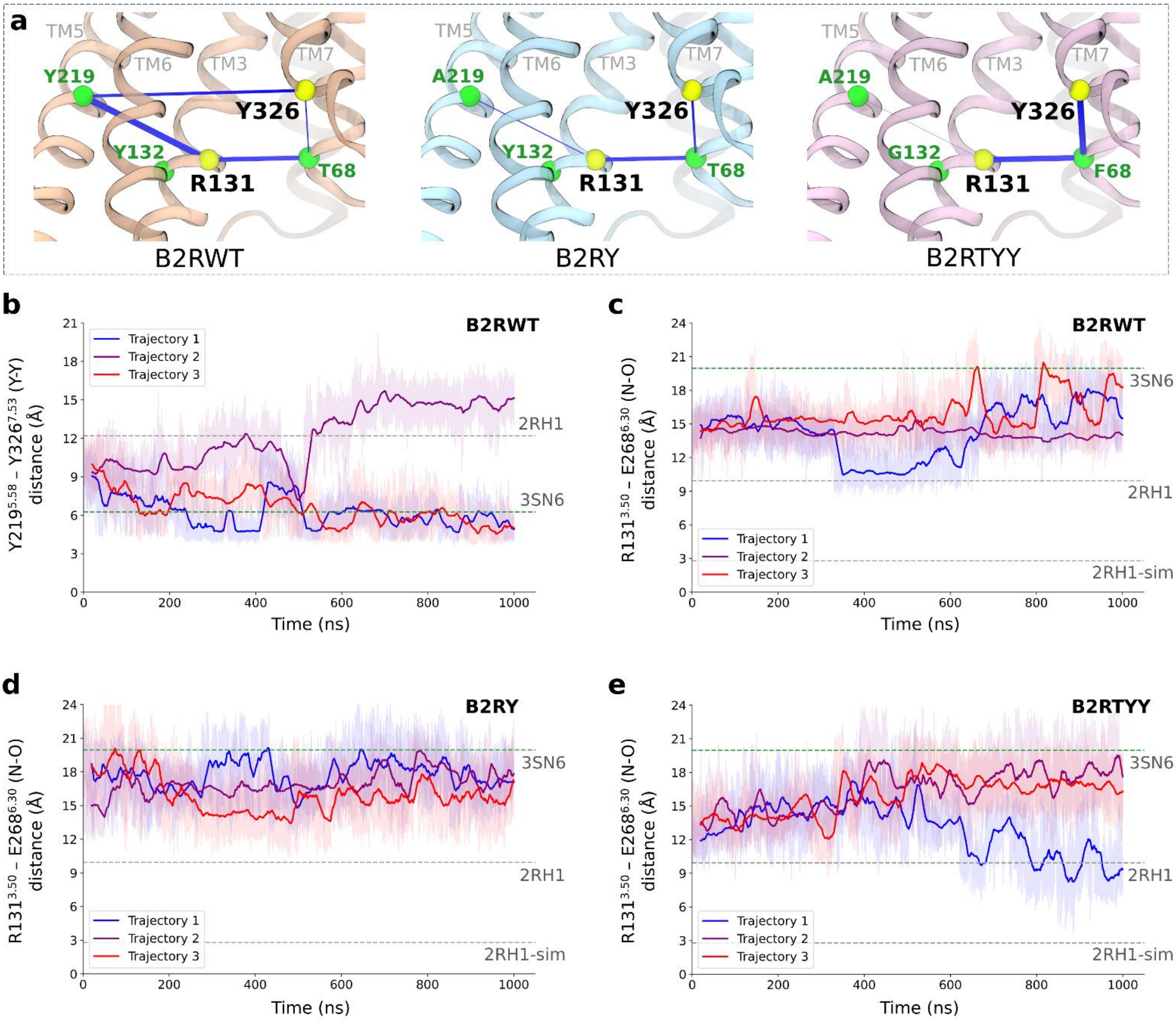


**Figure S10. (a)** Average nonbonding interaction energies of rotameric residues R131$^{3.50}$ and Y326$^{7.53}$ (yellow spheres) with mutated residues (green spheres). The numerical values of these interaction energies in kcal/mol are (i) B2RWT; R131-T68: -1.5, R131-Y219: -2, Y326-T68: -0.4, Y326-Y219: -1, (ii) B2RY; R131-T68: -1.3, R131-A219: -0.3, Y326-T68: -0.7, and (iii) B2RTYY; R131 F68: -1.8, R131 Y219: -0.1, Y326 T68: -2. **(b)** Trajectory-wise Y-Y distance, defined as the distance between $C_\zeta$ atoms of Y219$^{5.58}$ and Y326$^{7.53}$, indicates the formation of a water-mediated hydrogen bond,[27] (as observed in the active state structure - PDB ID: 3SN6) for the majority of the simulation time in the case of wild-type system. **(c-e)** Trajectory-wise N–O distance (the minimum distance between a guanidinium nitrogen atom of R131$^{3.50}$ and a carboxylate oxygen atom of E268$^{6.30}$) for all systems indicates a salt-bridge between these residues is not formed in our simulation. Smoothened lines are moving averages. Horizontal lines in green correspond to the values in PDB ID: 3SN6 (active) and grey in PDB ID: 2RH1 (inactive). 2RH1-sim indicates the value obtained for N-O distance in previous simulations[28] of the inactive state corresponding to the formation of a salt bridge between R131$^{3.50}$ and E268$^{6.30}$.

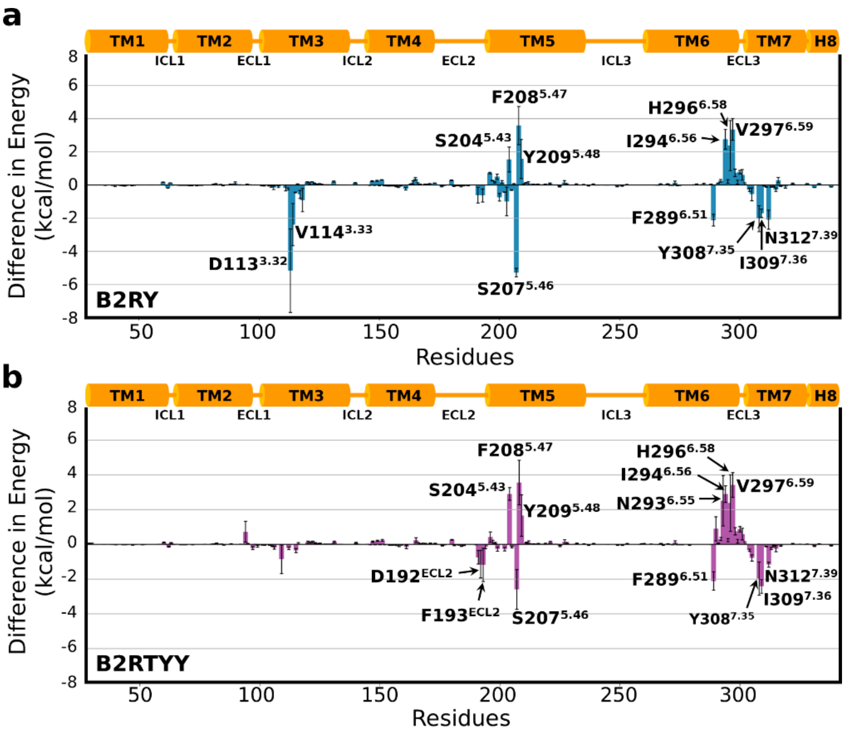


**Figure S11. (a)** The difference in nonbonding interaction energies ($\Delta E$) with agonist BI-167107 between **(a)** B2RY and B2RWT as well as **(b)** B2RTYY and B2RWT; residues with $|\Delta E| \geq 1$ kcal/mol are marked. The standard error of means from triplicate simulations are shown as error bars.

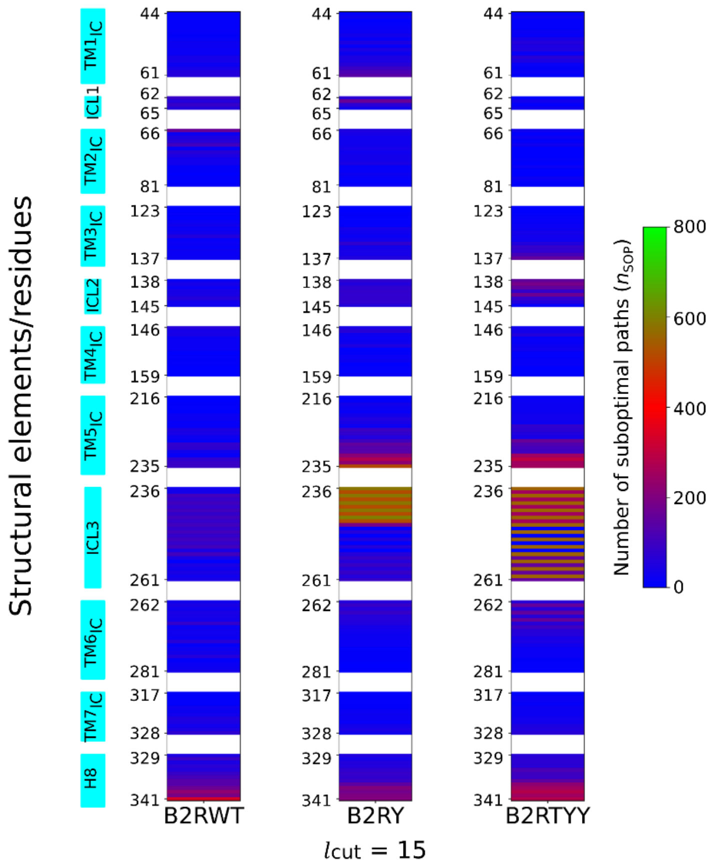


**Figure S12. (a)** Heatmap showing the number of suboptimal paths ($n_{SOP}$), from the agonist BI-167107 to the intracellular (IC) residues (residues at the cytosolic half of the receptor that are at least 10 Å apart from the agonist) for $l_{cut}$ = 15

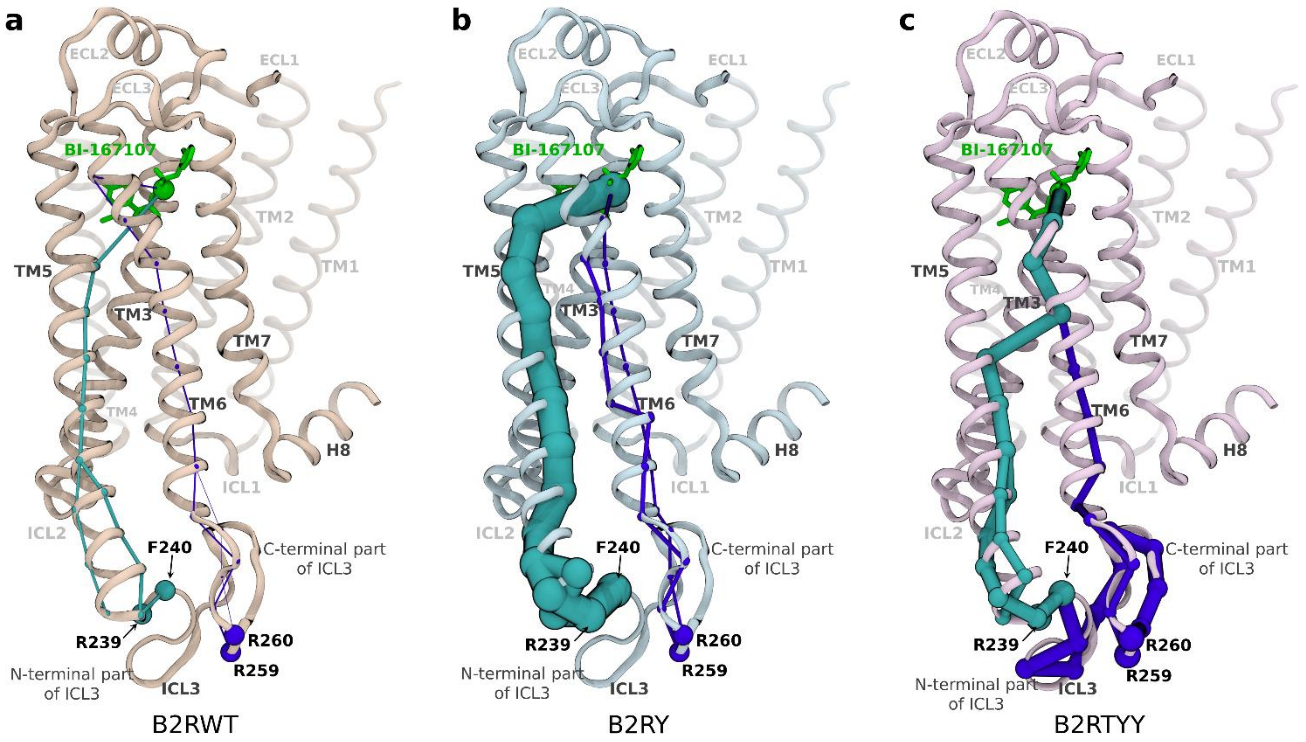


**Figure S13. (a-c)** Representations of allosteric communication paths (shortest paths with thickness proportional to $n_{SOP}$) from the agonist (green) to representative residues in the N-terminal (cyan) and C-terminal (violet) ends of ICL3.

### Supplementary tables

**Table S1.** GaMD simulation summary of the wild type (B2RWT), single mutant (B2RY), and triple mutant (B2RTYY) $\beta_2$AR systems. The average and standard deviation (σ) of total boost potentials ($\Delta V$) are tabulated.

| Simulation ID | Length (ns) | $\Delta V$ (kcal/mol) | |
|---|---|---|---|
| | | Avg | σ |
| **B2RWT** | | | |
| SIM1 | 1100 | 15.31 | 4.40 |
| SIM2 | 1100 | 15.28 | 4.40 |
| SIM3 | 1100 | 15.29 | 4.39 |
| **B2RY** | | | |
| SIM1 | 1100 | 14.96 | 4.36 |
| SIM2 | 1100 | 14.89 | 4.35 |
| SIM3 | 1100 | 15.18 | 4.39 |
| **B2RTYY** | | | |
| SIM1 | 1100 | 14.93 | 4.36 |
| SIM2 | 1100 | 14.77 | 4.32 |
| SIM3 | 1100 | 14.87 | 4.34 |

**Table S2.** Performance of metrics for all machine learning models implemented.

| **Classifying ML model** | **B2RWT** | | **B2RY** | | **B2RTYY** | |
|---|---|---|---|---|---|---|
| | Accuracy | Log loss | Accuracy | Log loss | Accuracy | Log loss |
| Random forest | 0.9962 | 0.0376 | 0.9953 | 0.0611 | 0.9912 | 0.0675 |
| Gradient Boosting | 0.9860 | 0.0606 | 0.9778 | 0.0921 | 0.9748 | 0.0915 |
| XGBoost | 0.9983 | 0.0054 | 0.9980 | 0.0068 | 0.9965 | 0.0112 |
| AutoGluon | 0.9983 | 0.0081 | 0.9986 | 0.0054 | 0.9978 | 0.0092 |

**Table S3.** Number of suboptimal paths ($n_{\text{SOP}}$) between the agonist BI-167107 and the residues in the intracellular halves of TM1-TM7, ICL1, ICL2, ICL3, and H8 that are at least 10Å apart from the agonist for $l_{\text{cut}} = 10$ and 15. Residues considered for depicting allosteric communication paths in Figures 6 and S13 are highlighted.

| Res ID | $n_{\text{SOP}}$ | | | | | | Res ID | $n_{\text{SOP}}$ | | | | | |
|---|---|---|---|---|---|---|---|---|---|---|---|---|---|
| | **B2RWT** | | **B2RY** | | **B2RTYY** | | | **B2RWT** | | **B2RY** | | **B2RTYY** | |
| | $l_{\text{cut}} = 10$ | $l_{\text{cut}} = 15$ | $l_{\text{cut}} = 10$ | $l_{\text{cut}} = 15$ | $l_{\text{cut}} = 10$ | $l_{\text{cut}} = 15$ | | $l_{\text{cut}} = 10$ | $l_{\text{cut}} = 15$ | $l_{\text{cut}} = 10$ | $l_{\text{cut}} = 15$ | $l_{\text{cut}} = 10$ | $l_{\text{cut}} = 15$ |
| V44$^{1.43}$ | 4 | 7 | 2 | 4 | 2 | 5 | Q224$^{5.63}$ | 3 | 13 | 8 | 20 | 27 | 80 |
| L45$^{1.44}$ | 4 | 7 | 8 | 22 | 6 | 20 | E225$^{5.64}$ | 28 | 49 | 38 | 94 | 22 | 71 |
| A46$^{1.45}$ | 4 | 7 | 2 | 4 | 4 | 8 | **A226$^{5.65}$** | **17** | **40** | **30** | **80** | **13** | **53** |
| I47$^{1.46}$ | 5 | 8 | 3 | 9 | 3 | 7 | K227$^{5.66}$ | 7 | 22 | 24 | 50 | 10 | 38 |
| V48$^{1.47}$ | 5 | 9 | 6 | 14 | 2 | 7 | R228$^{5.67}$ | 3 | 13 | 42 | 103 | 44 | 164 |
| F49$^{1.48}$ | 6 | 13 | 12 | 18 | 7 | 16 | **Q229$^{5.68}$** | **30** | **80** | **55** | **150** | **28** | **128** |
| G50$^{1.49}$ | 6 | 12 | 12 | 34 | 7 | 15 | L230$^{5.69}$ | 19 | 74 | 54 | 132 | 23 | 109 |
| N51$^{1.50}$ | 3 | 13 | 10 | 14 | 16 | 39 | Q231$^{5.70}$ | 9 | 27 | 48 | 136 | 29 | 99 |
| V52$^{1.51}$ | 10 | 25 | 15 | 48 | 23 | 57 | K232$^{5.71}$ | 3 | 15 | 75 | 216 | 55 | 253 |
| L53$^{1.52}$ | 5 | 20 | 10 | 14 | 19 | 46 | I233$^{5.72}$ | 24 | 78 | 106 | 273 | 62 | 298 |
| V54$^{1.53}$ | 4 | 16 | 14 | 45 | 21 | 52 | D234$^{5.73}$ | 21 | 86 | 70 | 220 | 58 | 253 |
| I55$^{1.54}$ | 4 | 16 | 12 | 25 | 7 | 14 | K235$^{5.74}$ | 25 | 93 | 184 | 521 | 75 | 254 |
| T56$^{1.55}$ | 8 | 35 | 14 | 48 | 31 | 66 | S236$^{ICL3}$ | 10 | 29 | 211 | 582 | 108 | 550 |
| A57$^{1.56}$ | 9 | 36 | 14 | 46 | 14 | 44 | E237$^{ICL3}$ | 5 | 33 | 184 | 521 | 75 | 255 |
| I58$^{1.57}$ | 5 | 20 | 19 | 65 | 10 | 14 | G238$^{ICL3}$ | 26 | 106 | 211 | 582 | 108 | 556 |
| A59$^{1.58}$ | 11 | 36 | 29 | 92 | 3 | 12 | **R239$^{ICL3}$** | **20** | **83** | **184** | **521** | **75** | **255** |
| K60$^{1.59}$ | 14 | 55 | 38 | 120 | 13 | 18 | **F240$^{ICL3}$** | **26** | **106** | **211** | **582** | **108** | **556** |
| F61$^{1.60}$ | 11 | 58 | 41 | 139 | 3 | 7 | H241$^{ICL3}$ | 20 | 83 | 184 | 521 | 75 | 255 |
| **E62$^{ICL1}$** | **40** | **90** | **20** | **66** | **3** | **7** | V242$^{ICL3}$ | 26 | 106 | 211 | 582 | 108 | 556 |
| R63$^{ICL1}$ | 11 | 58 | 52 | 164 | 16 | 29 | Q243$^{ICL3}$ | 20 | 83 | 184 | 521 | 75 | 255 |
| L64$^{ICL1}$ | 40 | 87 | 20 | 66 | 3 | 7 | N244$^{ICL3}$ | 26 | 106 | 215 | 597 | 108 | 556 |
| Q65$^{ICL1}$ | 11 | 58 | 11 | 29 | 14 | 26 | L245$^{ICL3}$ | 20 | 83 | 184 | 521 | 75 | 255 |
| T66$^{2.37}$ | 51 | 165 | 20 | 48 | 14 | 27 | S246$^{ICL3}$ | 26 | 106 | 46 | 189 | 108 | 556 |
| V67$^{2.38}$ | 7 | 24 | 14 | 38 | 5 | 9 | Q247$^{ICL3}$ | 20 | 83 | 0 | 0 | 0 | 0 |
| T68$^{2.39}$ | 12 | 44 | 11 | 29 | 11 | 20 | V248$^{ICL3}$ | 26 | 106 | 22 | 58 | 108 | 556 |
| N69$^{2.40}$ | 24 | 60 | 11 | 26 | 3 | 7 | E249$^{ICL3}$ | 20 | 83 | 0 | 0 | 0 | 0 |
| Y70$^{2.41}$ | 45 | 106 | 11 | 29 | 14 | 26 | Q250$^{ICL3}$ | 26 | 106 | 22 | 58 | 108 | 556 |
| F71$^{2.42}$ | 5 | 14 | 11 | 36 | 3 | 6 | D251$^{ICL3}$ | 20 | 83 | 0 | 0 | 0 | 0 |
| I72$^{2.43}$ | 28 | 65 | 11 | 26 | 2 | 4 | G252$^{ICL3}$ | 26 | 106 | 22 | 58 | 108 | 556 |
| T73$^{2.44}$ | 22 | 44 | 20 | 34 | 4 | 4 | R253$^{ICL3}$ | 12 | 41 | 0 | 0 | 0 | 0 |
| S74$^{2.45}$ | 15 | 30 | 20 | 32 | 4 | 8 | T254$^{ICL3}$ | 26 | 106 | 22 | 58 | 108 | 556 |
| L75$^{2.46}$ | 16 | 31 | 19 | 42 | 1 | 1 | G255$^{ICL3}$ | 12 | 41 | 31 | 90 | 58 | 157 |
| A76$^{2.47}$ | 6 | 12 | 10 | 14 | 10 | 17 | H256$^{ICL3}$ | 0 | 0 | 22 | 58 | 108 | 556 |
| C77$^{2.48}$ | 6 | 12 | 10 | 14 | 1 | 3 | G257$^{ICL3}$ | 12 | 41 | 31 | 90 | 58 | 157 |
| A78$^{2.49}$ | 2 | 2 | 13 | 25 | 3 | 3 | L258$^{ICL3}$ | 3 | 29 | 22 | 58 | 108 | 556 |
| D79$^{2.50}$ | 6 | 12 | 10 | 14 | 8 | 21 | **R259$^{ICL3}$** | **12** | **41** | **31** | **90** | **58** | **157** |
| L80$^{2.51}$ | 2 | 4 | 3 | 4 | 6 | 9 | **R260$^{ICL3}$** | **3** | **29** | **22** | **58** | **108** | **556** |
| V81$^{2.52}$ | 6 | 12 | 10 | 15 | 4 | 6 | S261$^{ICL3}$ | 12 | 41 | 31 | 90 | 58 | 157 |
| T123$^{3.42}$ | 5 | 6 | 3 | 3 | 3 | 4 | S262$^{6.24}$ | 3 | 29 | 22 | 58 | 24 | 64 |
| L124$^{3.43}$ | 5 | 6 | 2 | 4 | 4 | 5 | K263$^{6.25}$ | 12 | 41 | 31 | 90 | 58 | 157 |
| C125$^{3.44}$ | 5 | 8 | 4 | 11 | 2 | 7 | F264$^{6.26}$ | 3 | 29 | 22 | 58 | 24 | 64 |
| V126$^{3.45}$ | 1 | 5 | 5 | 7 | 10 | 13 | C265$^{6.27}$ | 12 | 41 | 31 | 90 | 58 | 157 |
| I127$^{3.46}$ | 13 | 17 | 5 | 7 | 8 | 13 | L266$^{6.28}$ | 14 | 36 | 22 | 58 | 24 | 64 |
| A128$^{3.47}$ | 21 | 31 | 5 | 9 | 7 | 16 | K267$^{6.29}$ | 11 | 32 | 23 | 58 | 58 | 157 |
| V129$^{3.48}$ | 9 | 17 | 16 | 36 | 8 | 21 | E268$^{6.30}$ | 36 | 79 | 22 | 51 | 24 | 64 |
| D130$^{3.49}$ | 7 | 26 | 13 | 26 | 20 | 36 | H269$^{6.31}$ | 2 | 18 | 16 | 37 | 44 | 103 |

| | | | | | | | | | | | | | |
|---|---|---|---|---|---|---|---|---|---|---|---|---|---|
| R131$^{3.50}$ | 36 | 56 | 8 | 17 | 15 | 30 | K270$^{6.32}$ | 10 | 28 | 12 | 30 | 24 | 64 |
| Y132$^{3.51}$ | 7 | 14 | 16 | 33 | 9 | 28 | A271$^{6.33}$ | 9 | 28 | 10 | 22 | 20 | 59 |
| F133$^{3.52}$ | 10 | 18 | 25 | 78 | 27 | 56 | L272$^{6.34}$ | 3 | 21 | 9 | 20 | 16 | 40 |
| **A134$^{3.53}$** | **8** | **15** | **16** | **34** | **41** | **82** | K273$^{6.35}$ | 25 | 63 | 10 | 18 | 23 | 43 |
| I135$^{3.54}$ | 7 | 19 | 16 | 40 | 27 | 76 | T274$^{6.36}$ | 2 | 11 | 8 | 16 | 13 | 30 |
| T136$^{3.55}$ | 7 | 18 | 16 | 37 | 42 | 111 | L275$^{6.37}$ | 9 | 23 | 3 | 7 | 5 | 17 |
| S137$^{3.56}$ | 10 | 20 | 25 | 56 | 56 | 156 | G276$^{6.38}$ | 15 | 33 | 6 | 10 | 9 | 18 |
| P138$^{ICL2}$ | 7 | 18 | 25 | 78 | 56 | 156 | I277$^{6.39}$ | 25 | 58 | 7 | 11 | 9 | 17 |
| F139$^{ICL2}$ | 18 | 40 | 25 | 56 | 63 | 185 | I278$^{6.40}$ | 2 | 7 | 7 | 12 | 6 | 12 |
| **K140$^{ICL2}$** | **10** | **20** | **25** | **78** | **56** | **156** | M279$^{6.41}$ | 7 | 16 | 3 | 4 | 3 | 8 |
| Y141$^{ICL2}$ | 13 | 38 | 35 | 81 | 41 | 83 | G280$^{6.42}$ | 10 | 23 | 4 | 6 | 4 | 7 |
| Q142$^{ICL2}$ | 21 | 43 | 25 | 78 | 57 | 169 | T281$^{6.43}$ | 14 | 32 | 5 | 8 | 3 | 8 |
| S143$^{ICL2}$ | 23 | 60 | 35 | 81 | 41 | 83 | V317$^{7.44}$ | 7 | 11 | 2 | 5 | 1 | 6 |
| L144$^{ICL2}$ | 14 | 31 | 25 | 78 | 27 | 56 | N318$^{7.45}$ | 7 | 12 | 5 | 10 | 7 | 17 |
| L145$^{ICL2}$ | 7 | 28 | 18 | 59 | 5 | 9 | S319$^{7.46}$ | 5 | 9 | 7 | 9 | 7 | 17 |
| **T146$^{4.38}$** | **23** | **57** | **20** | **44** | **6** | **29** | G320$^{7.47}$ | 2 | 8 | 5 | 7 | 4 | 14 |
| K147$^{4.39}$ | 18 | 36 | 28 | 63 | 10 | 34 | F321$^{7.48}$ | 14 | 28 | 4 | 11 | 6 | 19 |
| N148$^{4.40}$ | 26 | 38 | 3 | 22 | 17 | 54 | N322$^{7.49}$ | 11 | 25 | 6 | 17 | 8 | 21 |
| K149$^{4.41}$ | 5 | 16 | 7 | 16 | 2 | 13 | P323$^{7.50}$ | 15 | 30 | 11 | 25 | 8 | 25 |
| A150$^{4.42}$ | 7 | 17 | 11 | 23 | 4 | 13 | L324$^{7.51}$ | 25 | 53 | 7 | 19 | 11 | 29 |
| R151$^{4.43}$ | 12 | 18 | 17 | 48 | 6 | 22 | I325$^{7.52}$ | 22 | 45 | 4 | 13 | 11 | 31 |
| V152$^{4.44}$ | 12 | 17 | 3 | 6 | 9 | 28 | **Y326$^{7.53}$** | **13** | **29** | **11** | **34** | **19** | **50** |
| I153$^{4.45}$ | 3 | 6 | 6 | 12 | 1 | 7 | **C327$^{7.54}$** | **22** | **45** | **14** | **42** | **21** | **52** |
| I154$^{4.46}$ | 4 | 8 | 8 | 16 | 2 | 9 | R328$^{7.55}$ | 35 | 74 | 12 | 34 | 24 | 64 |
| L155$^{4.47}$ | 8 | 10 | 5 | 22 | 4 | 14 | **S329$^{8.47}$** | **22** | **45** | **16** | **53** | **24** | **63** |
| M156$^{4.48}$ | 8 | 8 | 3 | 4 | 5 | 15 | **P330$^{8.48}$** | **48** | **91** | **12** | **37** | **25** | **70** |
| V157$^{4.49}$ | 3 | 3 | 3 | 8 | 1 | 4 | **D331$^{8.49}$** | **22** | **46** | **16** | **55** | **24** | **70** |
| W158$^{4.50}$ | 3 | 4 | 2 | 7 | 2 | 5 | F332$^{8.50}$ | 35 | 67 | 17 | 56 | 23 | 68 |
| I159$^{4.51}$ | 4 | 4 | 11 | 16 | 6 | 10 | **R333$^{8.51}$** | **23** | **59** | **26** | **75** | **36** | **119** |
| V216$^{5.55}$ | 3 | 4 | 6 | 12 | 9 | 14 | I334$^{8.52}$ | 45 | 90 | 27 | 86 | 29 | 102 |
| F217$^{5.56}$ | 3 | 5 | 7 | 11 | 6 | 18 | A335$^{8.53}$ | 53 | 110 | 23 | 81 | 27 | 93 |
| V218$^{5.57}$ | 7 | 12 | 16 | 33 | 7 | 20 | F336$^{8.54}$ | 62 | 135 | 29 | 92 | 44 | 149 |
| Y219$^{5.58}$ | 6 | 8 | 13 | 25 | 7 | 20 | Q337$^{8.55}$ | 45 | 137 | 50 | 148 | 61 | 222 |
| S220$^{5.59}$ | 3 | 5 | 7 | 14 | 11 | 22 | E338$^{8.56}$ | 75 | 171 | 60 | 209 | 64 | 251 |
| R221$^{5.60}$ | 13 | 21 | 15 | 34 | 10 | 31 | L339$^{8.57}$ | 92 | 226 | 57 | 183 | 80 | 284 |
| **V222$^{5.61}$** | **10** | **19** | **24** | **50** | **10** | **29** | L340$^{8.58}$ | 73 | 192 | 56 | 199 | 64 | 251 |
| F223$^{5.62}$ | 8 | 15 | 20 | 40 | 8 | 25 | C341$^{8.59}$ | 114 | 337 | 61 | 205 | 80 | 284 |

**Table S4.** List of residues at the transducer (G protein/GRK/$\beta$-arrestin) interfaces of $\beta_2$AR. The residues at the G protein interface are obtained from the $\beta_2$AR-G$_s$ complex structure (PDB ID: 3SN6), and the residues at the $\beta$-arrestin/GRK interface are determined by aligning the crystal structure of the receptor (from the $\beta_2$AR-G$_s$ complex, PDB ID: 3SN6) with available $\beta$-arrestin/GRK complex structures of class A GPCRs; the $\beta_2$AR residues within 4 Å proximity of $\beta$-arrestin/GRK are identified as interface residues (see SI methods for details).

| G protein interface residues | GRK interface residues | $\beta$-arrestin interface residues |
|---|---|---|
| E62$^{ICL1}$, R63$^{ICL1}$, R131$^{3.50}$, A134$^{3.53}$, I135$^{3.54}$, T136$^{3.55}$, P138$^{ICL2}$, F139$^{ICL2}$, K140$^{ICL2}$, Y141$^{ICL2}$, Q142$^{ICL2}$, S143$^{ICL2}$, V222$^{5.61}$, E225$^{5.64}$, A226$^{5.65}$, R228$^{5.67}$, Q229$^{5.68}$, L230$^{5.69}$, K232$^{5.71}$, I233$^{5.72}$, K235$^{5.74}$, S236$^{5.75}$, R239$^{ICL3}$, K267$^{6.29}$, K270$^{6.32}$, A271$^{6.33}$, T274$^{6.36}$, L275$^{6.37}$ | A59$^{1.58}$, E62$^{ICL1}$, R63$^{ICL1}$, Q65$^{ICL1}$, T66$^{2.37}$, R131$^{3.50}$, F133$^{3.52}$, A134$^{3.53}$, I135$^{3.54}$, S137$^{3.56}$, F139$^{ICL2}$, K140$^{ICL2}$, Y141$^{ICL2}$, Q142$^{ICL2}$, S143$^{ICL2}$, L144$^{ICL2}$, T146$^{4.38}$, K147$^{4.39}$, N148$^{4.40}$, K149$^{4.41}$, R228$^{5.67}$, Q229$^{5.68}$, K232$^{5.71}$, I233$^{5.72}$, F264$^{6.26}$, K267$^{6.29}$, E268$^{6.30}$, K270$^{6.32}$, A271$^{6.33}$, T274$^{6.36}$, I325$^{7.52}$, Y326$^{7.53}$, S329$^{8.47}$, P330$^{8.48}$, D331$^{8.49}$, R333$^{8.51}$, I334$^{8.52}$ | R63$^{ICL1}$, Q65$^{ICL1}$, T66$^{2.37}$, V67$^{2.38}$, N69$^{2.40}$, I72$^{2.43}$, I127$^{3.46}$, R131$^{3.50}$, A134$^{3.53}$, I135$^{3.54}$, T136$^{3.55}$, S137$^{3.56}$, P138$^{ICL2}$, F139$^{ICL2}$, K140$^{ICL2}$, Y141$^{ICL2}$, Q142$^{ICL2}$, S143$^{ICL2}$, L144$^{ICL2}$, L145$^{ICL2}$, T146$^{4.38}$, K147$^{4.39}$ , E225$^{5.64}$, A226$^{5.65}$, Q229$^{5.68}$, K232$^{5.71}$, I233$^{5.72}$, K263$^{6.25}$, F264$^{6.26}$, L266$^{6.28}$, K267$^{6.29}$, E268$^{6.30}$ , K270$^{6.32}$, A271$^{6.33}$, T274$^{6.36}$, L275$^{6.37}$, I278$^{6.40}$, Y326$^{7.53}$, S329$^{8.47}$, P330$^{8.48}$, D331$^{8.49}$, F332$^{8.50}$ |
